# EZH2 Engages TGFβ Signaling to Promote Breast Cancer Bone Metastasis via Integrin β1-FAK Activation

**DOI:** 10.1101/2021.02.14.431151

**Authors:** Lin Zhang, Jingkun Qu, Yutao Qi, Yu-Wen Huang, Zhifen Zhou, Ping Li, Jun Yao, Beibei Huang, Shuxing Zhang, Dihua Yu

## Abstract

Bone metastasis is a frequent complication of breast cancer, occurring in about 50-70% of breast cancer patients with late-stage disease. The lack of effective therapy suggests that the precise molecular mechanisms underlying bone metastasis are still unclear. Enhancer of zeste homolog 2 (EZH2) is considered a breast cancer oncogene and its expression is correlated with metastasis of breast cancer, but its function in bone metastasis has not been well explored. Herein we report that EZH2 promotes osteolytic metastasis of breast cancer through regulating transforming growth factor beta (TGFβ) signaling, a key pathway in bone metastasis. Knocking down EZH2 decreases bone metastasis incidence and outgrowth *in vivo*. EZH2 induces cancer cell proliferation and osteoclast maturation, when breast cancer cells are co-cultured with osteoblasts and osteoclasts together *in vitro*. Mechanistically, EZH2 increases transcription of *ITGB1*, which encodes for integrin β1. Integrin β1 activates focal adhesion kinase (FAK), which phosphorylates TGFβ receptor type I (TGFβRI) at tyrosine 182, thus enhances the binding of TGFβRI to TGFβ receptor type II (TGFβRII), therefore activates Smad2 and increases parathyroid hormone-like hormone (PTHLH) expression. Clinically applicable FAK inhibitors but not EZH2 methyltransferase inhibitor effectively inhibits breast cancer bone metastasis *in vivo*. Overall, our data signify integrin β1-FAK as a new downstream effector of EZH2 in breast cancer cells, and EZH2-integrin β1-FAK axis cooperates with TGFβ signaling pathway to promote bone metastasis of breast cancer.

## Introduction

Breast cancer is the most commonly diagnosed cancer in female individuals worldwide^1^. About 50-70% of breast cancer patients with late-stage disease develop bone metastases that cause skeletal-related events, including pain, pathological fractures, spinal cord compression, hypercalcemia, and other complications^2^. The treatments for bone metastasis are limited and merely palliative; standard antiresorptive agents, chemotherapy, and radiotherapy can delay or lessen skeletal-related events, but they cannot cure bone metastasis^3^. Exploring the molecular mechanism of bone metastasis comprehensively may provide new therapeutic strategies for patients with bone metastasis. Breast cancer bone metastasis frequently induces osteolytic lesions, which lead to massive bone resorption and bone fractures^4^. Osteolytic bone resorption causes secretion of several growth factors, including transforming growth factor beta (TGFβ). Bone metastasis is incited by “the vicious cycle”, which designates the feed-forward cycle among cancer cells, osteoblasts, and osteoclasts in promoting both uncontrolled tumor growth and osteoclast activity^4–6^.

TGFβ plays dual roles in cancer initiation and progression: it works as a tumor suppressor in premalignant cells but induces breast cancer metastasis by enhancing epithelial-mesenchymal transition, angiogenesis, and immunosuppression^7,8^. Studies have well established that TGFβ is a predominant cytokine driving the feed-forward vicious cycle to promote metastatic cancer cell growth in bones^9^. In canonical TGFβ signaling, active TGFβ binds to its receptor, TGFβ receptor type II (TGFβRII), which binds and actives TGFβ receptor type I (TGFβRI) on the cell membrane. TGFβRI phosphorylates downstream signaling molecules Smad2/3, which form a complex with Smad4; the Smad2/3/4 complex is then translocated to the nucleus. The nuclear Smad2/3/4 complex works as a transcription factor to turn on the transcription of target genes^10,11^. Noncanonical TGFβ signaling works as a Smad-independent pathway through activation of p38 mitogen-activated protein kinase (MAPK), extracellular signal-regulated kinase (ERK), c-Jun N-terminal kinase (JNK), or phosphoinositide 3-kinase (PI3K)/AKT signaling^12^.

EZH2 is a histone methyltransferase that serves as an enzymatic subunit of the polycomb repressive complex 2^13^. It regulates gene expression through trimethylation of histone H3 at lysine (K) 27 (H3K27me3) or as a transcription co-factor^13^. Overexpression of EZH2 is correlated with metastasis of solid tumors such as prostate and breast cancers^14,15^ and is considered a prognostic biomarker of metastasis risk in women with early-stage hereditary breast cancer^16^. It is reported that EZH2 was highly expressed in tissues of renal cell carcinoma obtained from patients who had bone metastases^17^, suggesting that EZH2 promotes cancer cell bone metastasis. However, the function of EZH2 in the vicious cycle of breast cancer bone metastasis is unknown.

Here we found that depletion of EZH2 blocked breast cancer bone metastasis *in vivo*. Under TGFβ stimulation, EZH2 increased the level of pS465/467-Smad2 and the expression of parathyroid hormone-like hormone (PTHLH, also named parathyroid hormone-related protein, PTHRP), two key effectors of the canonical TGFβ pathway. Mechanistically, EZH2 increases the transcription of integrin β1-encoding *ITGB1* that activates downstream effector, focal adhesion kinase (FAK). Activated FAK phosphorylates TGFβRI and enhances the binding of TGFβRI to TGFβRII to active the TGFβ pathway. Our study revealed the cooperation between EZH2 and TGFβ signaling in promoting bone metastasis of breast cancer through a methyltransferase-independent mechanism, and demonstrated that targeting FAK may be an effective strategy for treatment of EZH2-induced breast cancer bone metastasis.

## Results

### EZH2 promotes breast cancer bone metastasis, which cannot be blocked by an EZH2 methyltransferase inhibitor

To explore the function of EZH2 in bone metastasis of breast cancer, we transfected either EZH2 shRNA or control shRNA into the MDA-MB-231 bone-seeking 231-1566 cell subline that expresses GFP and luciferase^7^ to generate the EZH2-knockdown cell lines 1566.shEZH2 and its control cell line 1566.shScr, respectively (Supplementary Fig. 1a). The sublines 1566.shEZH2 and 1566.shScr were injected, separately, into the left ventricles of nude mice. Mice injected with 1566.shEZH cells had significantly longer bone metastasis-free survival (*P* = 0.0047) and overall survival (*P* = 0.0024) than did mice injected with 1566.shScr cells (Fig. 1a, b). Bioluminescence imaging (BLI), X-ray imaging, and hematoxylin and eosin (H&E) staining of bone lesions all showed that mice injected with 1566.shEZH cells had fewer bone metastases than did mice injected with 1566.shScr cells on the same day post-injection (Fig. 1b). Using the CRISPR/CAS9 system, EZH2-knockout MDA-MB-231 cell subclones (231.KO) and their control clone (231.sgCtrl) were also generated (Supplementary Fig. 1b), and one of the 231.KO subclones (231.KO#1) were stably re-expressed with wild-type EZH2 (231.KO#1.EZH2) or a pLenti control vector (231.KO#1.pLenti) (Supplementary Fig. 1c). The derived sublines 231.KO#1.EZH2 and 231.KO#1.pLenti were intracardially injected individually into nude mice. Mice injected with 231.KO#1.EZH2 cells were treated with a vehicle or GSK126, a potent small-molecule EZH2 methyltransferase inhibitor, whereas the control mice injected with 231.KO#1.pLenti were only treated with a vehicle. The results showed that the vehicle-treated 231.KO#1.EZH2 group had poorer bone metastasis-free survival rates than did the control 231.KO#1.pLenti group (Fig. 1c). Unexpectedly, the GSK126-treated 231.KO#1.EZH2 group had similar metastasis-free survival rate to that in the vehicle-treated 231.KO#1.EZH2 group (Fig. 1c, d). The data indicated that EZH2 overexpression increased the incidence of bone metastasis, which cannot be deterred by inhibiting EZH2 methyltransferase function with GSK126. The above loss- and gain-of EZH2 function *in vivo* bone metastasis experiments demonstrated that EZH2 promoted breast cancer bone metastasis and that EZH2’s effect on bone metastasis is likely methyltransferase-independent.

**Figure 1.**
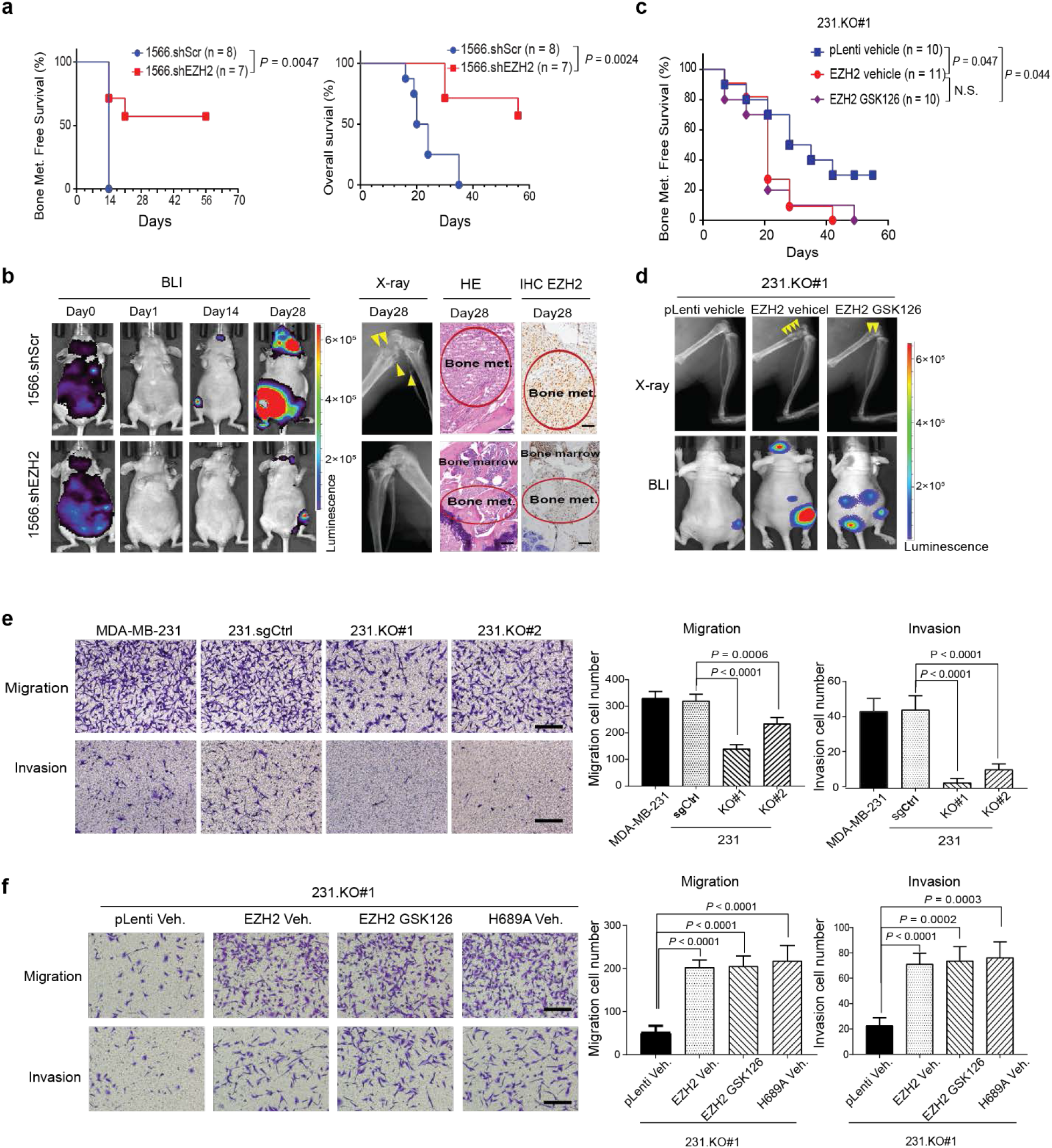
EZH2 promotes breast cancer bone metastasis, which cannot be blocked by an EZH2 methyltransferase inhibitor. **a,** Kaplan-Meier curves showing bone metastasis-free survival and overall survival rates in mice intracardially injected with 1 × 10^5^ 1566.shCtrl (*n* = 8) or 1566.shEZH2 (*n* = 7) cells. Log-rank test. **b,** Representative bioluminescence (BLI), X-ray, and hematoxylin and eosin (HE)-stained images of bone-metastatic lesions in the 2 subgroups described in **a** obtained at the indicated time points. The arrows indicate osteolytic bone lesions in X-ray images. EZH2 expression is shown by IHC staining. Scale bars, 50 μm. **c,** Kaplan-Meier curves showing bone metastasis-free survival rates in mice intracardially injected with 1 × 10^5^ 231.KO#1.pLenti cells and given treatment with a vehicle (231.KO#1.pLenti vehicle*, n* = 10) or 1 × 10^5^ 231.KO#1.EZH2 cells and given treatment with a vehicle (231.KO#1.EZH2 vehicle*, n* = 11) or GSK126 (231.KO#1.EZH2 GSK126; *n* = 10; 150 mg/kg GSK126 per mouse, i.p. injection three times a week). Log-rank test. **d,** Representative X-ray and bioluminescence (BLI) images of bone-metastatic lesions in the 3 subgroups described in **c**. **e,** Representative images and quantification of invading and migrating MDA-MB-231, 231.sgCtrl., 231.KO#1, and 231.KO#2 cells. Scale bars, 100 μm. Data are presented as means ± S.D. (*t*-test). **f,** Representative images and quantification of invading and migrating 231.KO#1.pLenti. cells treated with vehicle, 231.KO#1.EZH2 cells treated with vehicle, 231.KO#1.EZH2 cells treated with 2 μM GSK126, and 231.KO#1.H689A cells treated with vehicle. Scale bars, 100 μm. Data are presented as means ± S.D. (*t*-test).

To explore the mechanism of EZH2 promotion of bone metastasis, we first compared the proliferation, migration, and invasion ability of high and low EZH2-expressing MDA-MB-231 cell sublines. High EZH2-expressing cells (MDA-MB-231 and 231.sgCtrl) and low EZH2-expressing cells (231.KO #1 and #2) had similar rates of proliferation in two-dimensional cell culture as measured using MMT assays (Supplementary Fig. 1d). However, high EZH2-expressing cells had greater migration and invasion ability *in vitro* than did low EZH2-expressing cells (Fig. 1e). GSK126 treatment didn’t change cell proliferation, migration, or invasion of high EZH2-expressing MDA-MB-231 cells compared to vehicle treatment (Supplementary Fig. 1e-h). In addition to the 231.KO#1.pLenti, 231.KO#1.EZH2 sublines, we also re-expressed in 231.KO#1 cells an EZH2-methyltransferase-dead mutant EZH2-H689A to generate the stable 231.KO#1.H689A subline (Supplementary Fig. 1c). Re-expression of wild-type EZH2 (231.KO#1.EZH2) greatly increased cell migration and invasion compared to the control 231.KO#1.pLenti cells (Fig. 1f). And re-expression of methyltransferase-dead mutant EZH2 H689A in 231.KO#1 (231.KO#1.H689A) also promoted migration, or invasion just like that of re-expression of wild-type EZH2 (231.KO#1.EZH2) (Fig. 1f). Similarly, GSK126-treatment didn’t have any inhibitory effect on migration, or invasion of 231.KO#1.EZH2 cells (Fig. 1f). These data suggested that EZH2 promoted MDA-MB-231 cell migration and invasion but did not change cell proliferation in two-dimensional cell culture. In addition, blocking EZH2’s histone methyltransferase function, either genetically or with EZH2 methyltransferase inhibitor GSK126 did not inhibit the migration or invasive ability induced by wild-type EZH2.

To expand the investigation of EZH2’s effect on bone metastasis, we also established CRISPR/CAS9-mediated EZH2-knockout subclones in 4T1 mouse mammary tumor cells (4T1.KO #1 and #2) (Supplementary Fig. 1i) and examined their proliferation, migration, and invasion compared with those of the control 4T1 cells. Knocking out EZH2 inhibited 4T1 cell migration and invasion but did not have a dramatic effect on cell proliferation (Supplementary Fig. 1j, k). GSK126-based treatment of 4T1 cells did not result in different proliferation, invasion, or migration from untreated 4T1 cells (Supplementary Fig. 1j-l). Clearly, our data from both MBA-MD-231 and 4T1 cells showed that EZH2 promoted cancer cell migration and invasion, but this function is unlikely dependent on EZH2’s methyltransferase activity.

### EZH2 regulates the vicious cycle of breast cancer bone metastasis

The colonization and growth of cancer cells in bone marrow is key to bone metastasis formation^6^. Thus, we explored the role of EZH2 in promoting metastatic breast cancer outgrowth in the bone. To mimic the vicious cycle of breast cancer bone metastasis microenvironment, we co-cultured breast cancer cells with RAW264.7 preosteoclasts and MC3T3 osteoblasts (triple co-culture) under TGFβ treatment (5 ng/mL) (Fig. 2a). The MDA-MB-231, 231.sgCtrl, 231.KO#1, and 231.KO#2 cells were pre-transfected with GFP expression vector for easy detection and quantification by flow cytometry (Supplementary Fig. 2a, b). Mature osteoclasts are detected by TRAP staining as round giant cells with three or more nuclei^18^ and they induce osteolysis to release TGFβ, which activates the vicious cycle of breast cancer bone metastasis^9^. Six days in triple co-culture, the EZH2-knockout 231.KO#1 and 231.KO#2 cells showed significantly inhibited cell growth than MDA-MB-231 and 231.sgCtrl cells (Fig. 2b and Supplementary Fig. 2b). Also, the RAW264.7 preosteoclasts that differentiated into mature osteoclasts were significantly less in co-culture with EZH2-knockout cells (231.KO#1 and 231.KO#2) than with MDA-MB-231 or 231.sgCtrl cells (Fig. 2c). When MDA-MB-231 cells were treated with 2 μM GSK126 or a vehicle (dimethyl sulfoxide, DMSO) in triple co-culture, GSK126 did not inhibit cancer cell proliferation or osteoclast cell maturation (Supplementary Fig. 2c, d). To further test EZH2 methyltransferase function in vicious cycle of breast cancer bone metastasis, 231.KO#1.pLenti, 231.KO#1.EZH2, and 231.KO#1.H689A cells were compared in triple co-culture. Similar to the wild-type EZH2-re-expressing 231.KO#1.EZH2 cells, 231.KO#1.H689A cells that re-expressed EZH2 methyltransferase-dead mutant H689A had enhanced tumor cell proliferation and osteoclast maturation compared to 231.KO#1.pLenti cells (Fig. 2d, e). In addition, EZH2-reexpressing 231.KO#1.EZH2 cells with or without GSK126 treatment showed similarly increased tumor cell proliferation and osteoclast maturation compared to 231.KO#1.pLenti cells (Fig. 2d, e). Likewise, we performed triple co-culture experiments with 4T1 cells and EZH2-knockout 4T1 cell sublines (4T1.KO #1 and #2) as well as treating 4T1 cells with GSK126, and had consistent findings as those from the MDA-MB-231 cell sublines. Mainly, i) high EZH2-expressing cells (4T1) had a growth advantage and induced osteoclasts maturation more than did EZH2-knockout cells; ii) GSK126 did not inhibit 4T1 cell proliferation or RAW264.7 preosteoclasts maturation in the triple co-culture (Supplementary Fig. 2e, f). Together, data from both cell models indicate that EZH2 promoted vicious cycle of breast cancer bone metastasis, which cannot be blocked by treatment with the EZH2 methyltransferase inhibitor GSK126.

**Figure 2.**
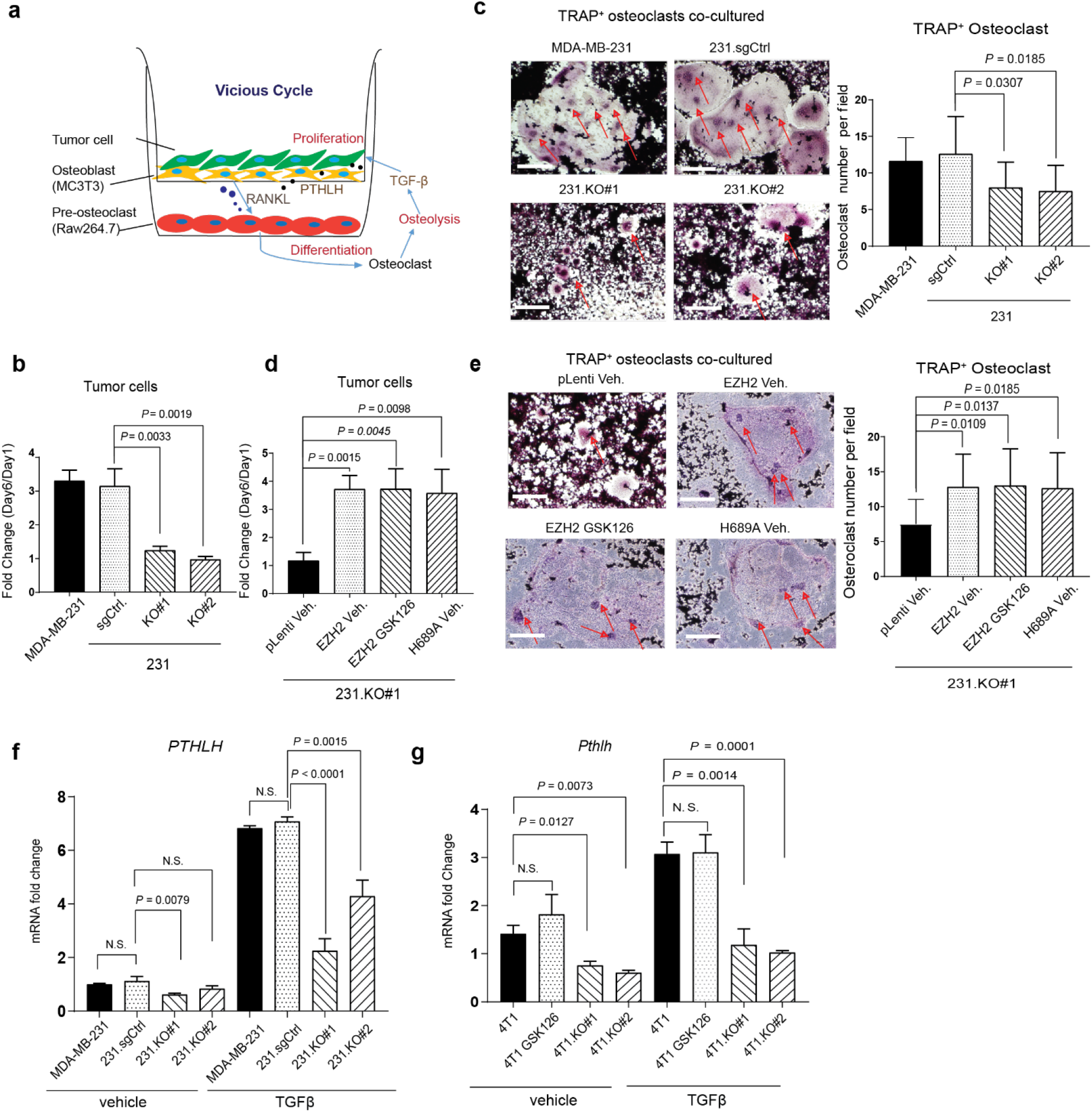
EZH2 regulates the vicious cycle of breast cancer bone metastasis. **a,** Model of triple co-culture of breast cancer cells with preosteoclasts and osetoblasts. Murine RAW 264.7 preosteoclasts were seeded into the wells of six-well plates. GFP-labeled breast cancer cells and MC3T3 osteoblasts were seeded into Millicell Hanging Cell Culture Inserts (Millipore) in the six-well plates and treated with TGFβ (5 ng/mL). **b,** Quantification of MDA-MB-231, 231.sgCtrl., 231.KO#1, and 231.KO#2 cells after co-culture with osteoclasts and MC3T3 osteoblasts treated with TGFβ (5 ng/mL) for 6 days. Data are presented as means ± S.D. (*t*-test). **c,** Representative staining images and quantification of mature TRAP^+^ osteoclasts after culture with MC3T3 osteoblasts and the indicated cancer cells, and treatment of them with TGFβ (5 ng/mL) for 6 days. The arrows indicate multinuclear mature TRAP^+^ osteoclasts. Scale bars, 200 μm. Data are presented as means ± S.D. (*t*-test). **d,** Quantification of 231.KO#1.pLenti. cells treated with vehicle, 231.KO#1.EZH2 cells treated with vehicle, 231.KO#1.EZH2 cells treated with 2 μM GSK126, 231.KO#1.H689A cells treated with vehicle, after co-culture with osteoclasts and MC3T3 osteoblasts treated with TGFβ (5 ng/mL) for 6 days. Data are presented as means ± S.D. (*t*-test). **e,** Representative staining images and quantification of mature TRAP^+^ osteoclasts after co-culture with MC3T3 osteoblasts and the indicated cancer cells and treatment of them with TGFβ (5 ng/mL) for 6 days. Scale bars, 200 μm. Data are presented as means ± S.D. (*t*-test). **f,** qRT-PCR analysis of *PTHLH* mRNA expression in the indicated cells treated with a vehicle or TGFβ (5 ng/mL, 2 hours). Data are presented as means ± S.D. (*t*-test). N.S., not significant. **g,** qRT-PCR analysis of *Pthlh* mRNA expression in the indicated cells treated with a vehicle or TGFβ (5 ng/mL, 2 hours). Data are presented as means ± S.D. (*t*-test).

Parathyroid hormone-like hormone (PTHLH, also named PTHRP) is an essential mediator of breast cancer bone metastasis, and metastatic cancer cells secrete PTHLH into the bone microenvironment to activate osteolysis^19^. Remarkably, knockout of EZH2 in both MDA-MB-231 and 4T1 cells reduced their *PTHLH* mRNA expression under TGFβ treatment as measured by qRT-PCR (Fig. 2f, g), whereas GSK126 treatment of 4T1 cells didn’t reduce *Pthlh* mRNA expression (Fig. 2g). Besides PTHLH, IL-8 is a cytokine that also regulates osteolysis in breast cancer bone metastasis^20^. Knockout of EZH2 in MDA-MB-231 and 4T1 cells also inhibited their *IL-8* mRNA expression, but GSK126 treatment did not change it (Supplementary Fig. 2g, h). These data indicated that EZH2 facilitates *PTHLH* and *IL-8* expressions, which can mediate the vicious cycle of breast cancer bone metastasis.

### EZH2 increases pS465/467-Smad2 and pY397-FAK levels in response to TGFβ stimulation

*PTHLH* is a well-known TGFβ downstream gene regulated by the p-Smad2/Gli2 transcription factor complex or p38 MAPK^7,21^. To further explore how EZH2 facilitates PTHLH and TGFβ signaling in breast cancer cells, we detected pS465/467-Smad2 and pT180/Y182-p38 MAPK levels in MDA-MB-231 sublines. In response to TGFβ stimulation, knockout of EZH2 in MDA-MB-231 cells inhibited pS465/467-Smad2 levels without significant changes of total Smad2, Smad3, or Smad4 protein expressions (Fig. 3a and Supplementary Fig. 3a). Knockout of EZH2 did not change the level of pT180/Y182-p38 MAPK, suggesting that EZH2 does not regulate TGFβ-p38 MAPK signaling (Fig. 3a). Knockdown of EZH2 by shRNAs (shEZH2#3 and shEZH#4) yielded similar results in MDA-MB-231 sublines (Fig. 3b). To examine whether EZH2-methyltransferase-function is involved in regulation of pS465/467-Smad2 levels, we measured pS465/467-SMAD2 levels in 231.KO#1 sublines that have re-expression of the control vector, wild-type EZH2, or H689A EZH2, in response to TGFβ (Fig. 3c). Like wild-type EZH2 re-expressing cells, H689A EZH2 re-expression also increased the level of pS465/467-Smad2 compared with the control vector (Fig. 3c). Furthermore, GSK126 treatment had no significant impact on increased pS465/467-Smad2 by TGFβ treatment (Supplementary Fig. 3b).

**Figure 3.**
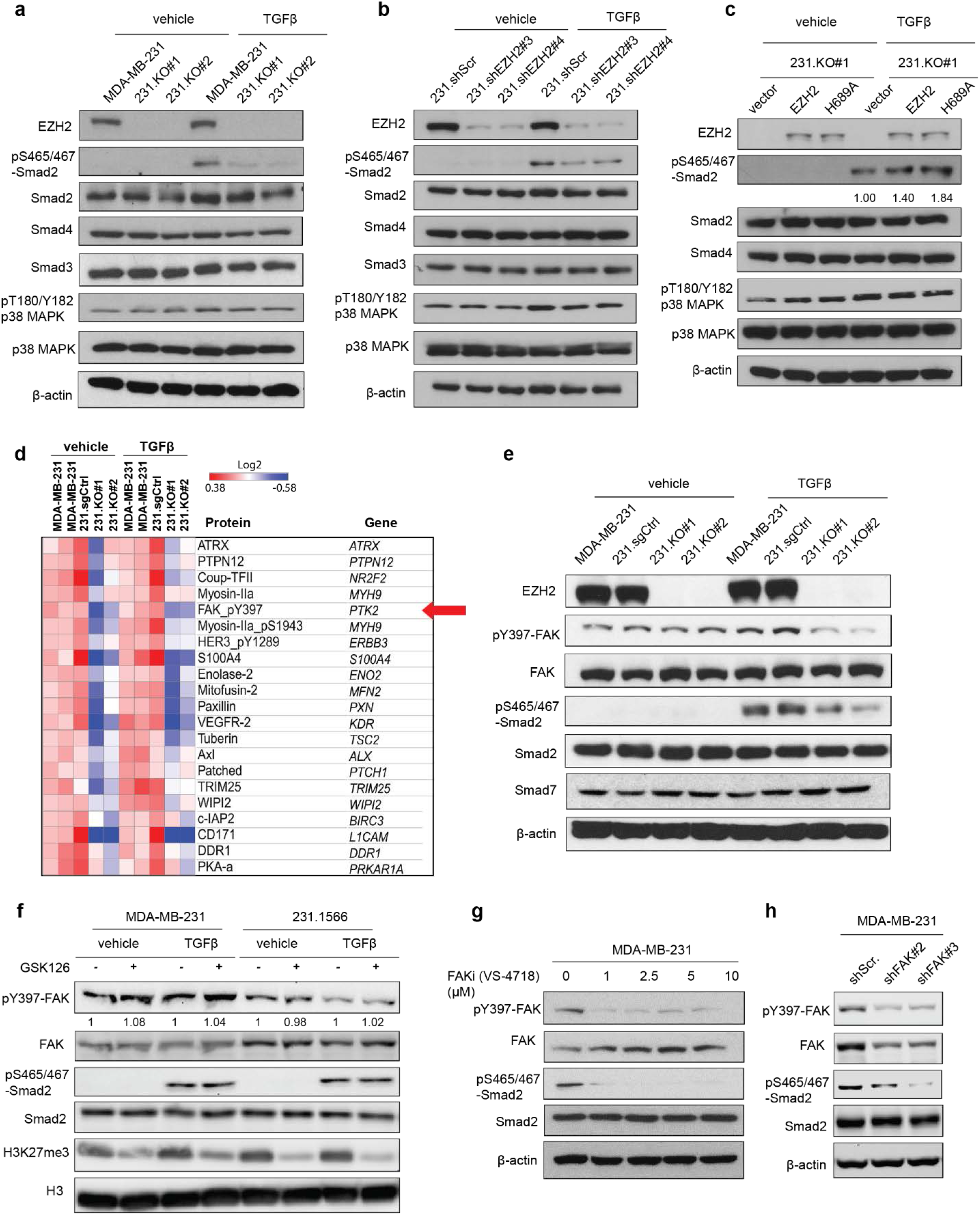
EZH2 increases pS465/467-Smad2 and pY397-FAK levels in response to TGFβ stimulation. **a,** Western blotting of the expression of the indicated proteins in MDA-MB-231, 231.KO#1, and 231.KO#2 cells treated with a vehicle or TGFβ (5 ng/mL) for 2 hours. **b,** Western blotting of the expression of the indicated proteins in 231.shScr, 231.shEZH2#3, and 231.shEZH2#4 cells treated with a vehicle or TGFβ (5 ng/mL) for 2 hours. **c,** Western blotting of the expression of the indicated proteins in 231.KO#1.vector, 231.KO#1.EZH2, and 231.KO#1.H689A cells treated with a vehicle or TGFβ (5 ng/mL) for 2 hours. **d,** RPPA analysis of MDA-MB-231, 231.sgCtrl, 231.KO#1, and 231.KO#2 cells treated with a vehicle or TGFβ (5 ng/mL) for 2 hours. The heat map shows the top downregulated proteins in 231.KO#1 and 231.KO#2 cells compared with MDA-MB-231 and 231.sgCtrl cells. **e,** Western blotting of the expression of the indicated proteins in MDA-MB-231, 231.sgCtrl, 231.KO#1, and 231.KO#2 cells treated with a vehicle or TGFβ (5 ng/mL, 2 hours). **f,** Western blotting of the expression of the indicated proteins in MDA-MB-231 or 231-1566 cells treated with a vehicle or GSK (2 μM, 24 hours) and then treated without or with TGFβ (5 ng/mL, 2 hours). **g,** Western blotting of the expression of the indicated proteins in MDA-MB-231 cells treated with the FAKi VS-4718 at different concentrations (0-10 μM) and with TGFβ (5 ng/mL) for 2 hours. **h,** Western blotting of the expression of the indicated proteins in 231.shScr, 231.shFAK#2, and 231.shFAK#3 cells treated with TGFβ (5 ng/mL) for 2 hours.

To gain insight into how EZH2 activates Smad2 signaling, we first measured the expressions of TGFβRI and TGFβRII, the TGFβ receptors upstream of pS465/467-Smad2, using flow cytometry or Western blotting, and detected no significant changes in EZH2-knockout sublines (Supplementary Fig. 3c, d). Next, we performed reverse-phase protein array (RPPA) to profile protein expression changes in MDA-MB-231 and 231.sgCtrl cells versus those in EZH2-knockout MDA-MB-231 cells (231.KO #1 and #2) with or without TGFβ treatment. RPPA revealed that 228 proteins were downregulated and 194 proteins were upregulated in the two EZH2-knockout cell lines compared to that in MDA-MB-231 and 231.sgCtrl cells (Table 1), and gene ontology molecular functional analysis showed that the dramatically downregulated and upregulated proteins were kinases, including tyrosine kinases (Fig. 3d, and Supplementary Fig. 3e, f). Notably, phosphorylation of tyrosine 397 on FAK (pY397-FAK) was significantly downregulated in the EZH2-knockout cells (Fig. 3d). FAK is a non-receptor tyrosine kinase that regulates the survival, proliferation, migration, and invasion of cancer cells and can impact on cancer development and progression^22,23^. Unlike the FAK upstream kinase Src, whose function in bone metastasis is well-documented^24–26^, the function of FAK in bone metastasis is unclear. We thus validated RPPA data by Western blotting, which showed that knocking out EZH2 in MDA-MB-231 and MCF7 cells inhibited pY397-FAK and pS465/467-Smad2 levels under TGFβ treatment (Fig. 3e and Supplementary Fig. 3g). GSK126 treatment of MDA-MB-231 and 231-1566 cells with or without TGFβ stimulation did not change pY397-FAK or pS465/467-Smad2 levels compare to vehicle treated cells (Fig. 3f). Collectively, these data suggested that EZH2 functions to activate FAK and Smad2 signaling under TGFβ stimulation.

To examine whether increased pY397-FAK is related to enhanced pS465/467-Smad2, we treated MDA-MB-231 cells with FAK inhibitors (FAKi, VS-4718 or VS-6063) at different concentrations (1-10 μM), followed with TGFβ treatment (2 μM, 2 hours), then detected pS465/467-Smad2. FAKi treatment dramatically blocked pS465/467-Smad2 level and reduced *PTHLH* mRNA expression even under TGFβ stimulation (Fig. 3g and Supplementary Fig. 3h, i). Additionally, knocked down FAK using shRNAs (shFAK#2 or shFAK#3) in MDA-MB-231 cells also inhibited pS465/467-Smad2 levels (Fig. 3h). In the bone-seeking MDA-MB-231 subline 231-1566, knocking down FAK alone with siRNAs (siFAK#1 or siFAK#2) did not change pS465/467-Smad2 levels (Supplementary Fig. 3j); however, doubly knocking down FAK and its closely related kinase PYK2 (or FAK2) with siRNAs (siPYK2#44, siPYK2#49, or siPYK2#50) dramatically reduced pS465/467-Smad2 levels (Supplementary Fig. 3j). Doubly knocking down FAK and PYK2 also inhibited *PTHLH* mRNA expression (Supplementary Fig. 3k). These data indicated that activation of FAK family kinases by EZH2 increased the phosphorylation of S465/467-Smad2 and activated the TGFβ/Smad2/PTHLH pathway in breast cancer cells.

### pY397-FAK induces TGFβRI tyrosine phosphorylation that enhances its binding to TGFβRII in response to TGFβ

We further explored the mechanism of how EZH2-mediated FAK activation induces pS465/467-Smad2. Smad7 can block the TGFβRI-induced pS465/467-Smad2^27^, but knocking down EZH2 did not change Smad7 expression (Fig. 3e). Thus, we assessed whether FAK regulates TGFβRI and TGFβRII expressions in MDA-MB-231 cells. Knocking down FAK did not change the protein expressions of TGFβRI or TGFβRII (Supplementary Fig. 4a, b). Next, we tested whether FAK can bind to Smad2 to phosphorylate Smad2 by immunoprecipitation (IP) of FAK followed with Western blotting of Smad2, which didn’t show detectable binding (Supplementary Fig. 4c). Surprisingly, FAK IP brought down TGFβRI, not TGFβRII, with or without TGFβ exposure (Fig. 4a). Additionally, blocking FAK kinase activity by FAKi VS-4718 treatment reduced the binding of FAK to TGFβRI (Fig. 4b). Since TGFβ treatment induces TGFβRI binding to TGFβRII and, consequently, increased pS465/467-Smad2 levels (Supplementary Fig. 4d), we postulated that pY397-FAK may phosphorylate TGFβRI that increases the binding affinity of TGFβRI for TGFβRII under TGFβ stimulation, leading to activation of Smad2 signaling. To test this, MDA-MB-231 cells were treated by the FAKi VS-4718 or had FAK knockdown by shRNAs (shFAK#21 or shFAK#3), and were treated with TGFβ. After collecting cell lysates, we performed TGFβRI IP followed by Western blotting of TGFβRII, which showed that FAK inhibition dramatically reduced the binding of TGFβRI to TGFβRII in response to TGFβ stimulation (Fig. 4c and Supplementary Fig. 4e, f). To explore whether FAK tyrosine kinase can phosphorylate TGFβRI, we performed IP to pull down TGFβRI from MDA-MB-231 cells (231.TGFβRI) or HEK 293FT cells transfected with exogenous FLAG-tagged wild-type TGFβRI (293FT.TGFβRI), then Western blotting with anti-phospho-tyrosine antibodies. We detected tyrosine phosphorylation on TGFβRI (Supplementary Fig. 4g, h), which are reduced by FAKi VS-6063 treatment (Fig. 4d). The data indicated that activated FAK can induce tyrosine phosphorylation of TGFβRI. To locate the site of tyrosine phosphorylation on TGFβRI, we performed mass spectrometric analysis which identified a novel TGFβRI phosphorylation site at tyrosine 182 (pY182), which is located in the glycine and serine residues enriched-domain (GS domain)^28^ (Fig. 4e). It was reported that after TGFβ binding, activated TGFβRII phosphorylates TGFβRI in the GS domain and thereby activates TGFβRI kinase function and the transduction of TGFβ signals^29^. And the newly identified TGFβRI phosphorylation site of pY182 at the “YDMT” sequence fits the Src Homology 2 (SH2) motif “YXXT” that FAK can bind to^30,31^ (Fig. 4e).

**Figure 4.**
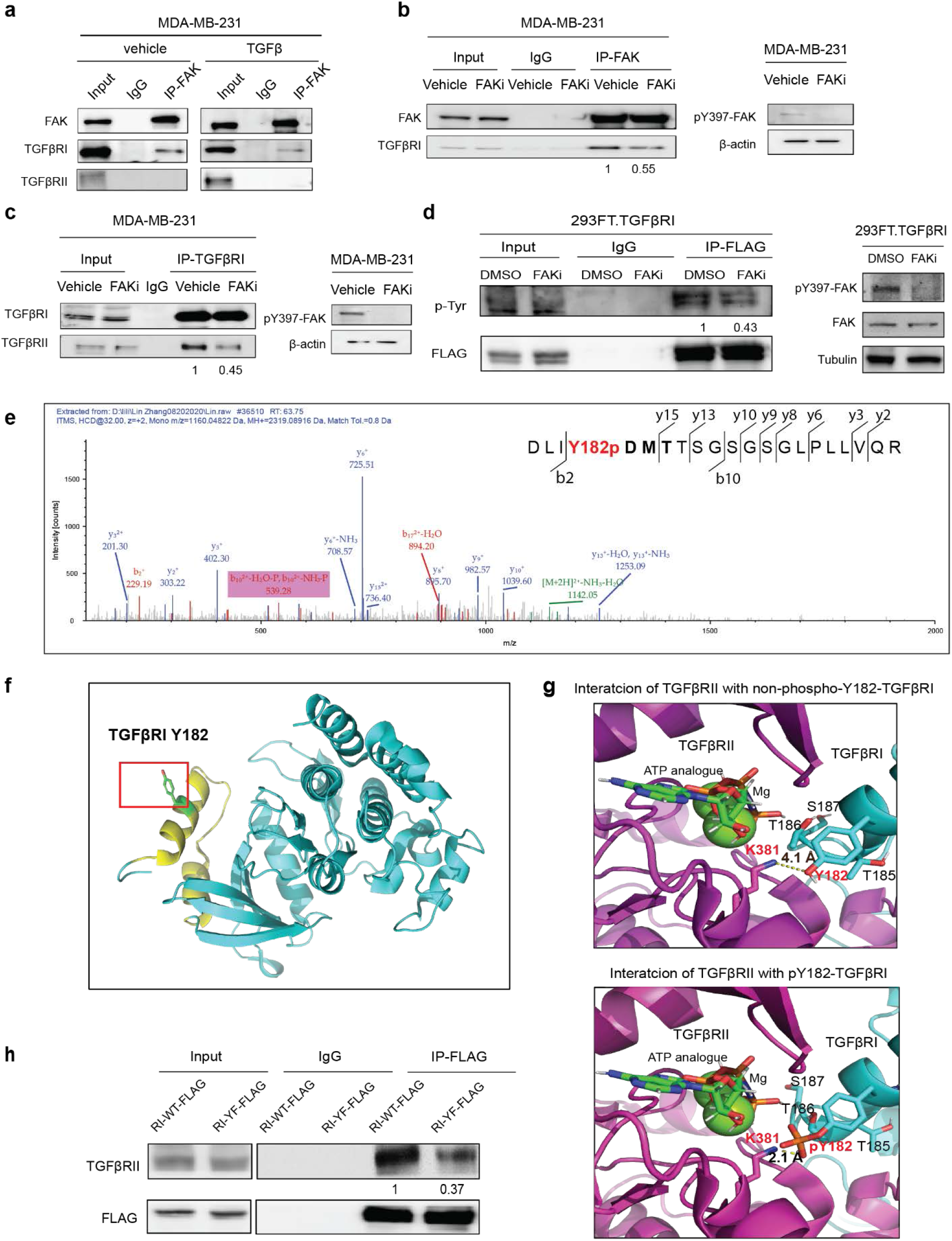
pY397-FAK induces TGFβRI tyrosine phosphorylation that enhances its binding to TGFβRII in response to TGFβ. **a,** IP of FAK from the lysis of MDA-MB-231 cells treated with a vehicle or TGFβ (5 ng/mL) for 2 hours followed by Western blotting for FAK, TGFβRI, and TGFβRII. **b,** IP of FAK from the lysis of MDA-MB-231 cells treated with a vehicle or the FAKi VS-4718 (1 μM) for 24 hours followed by Western blotting for FAK and TGFβRI. **c,** IP of TGFβRI from the lysis of MDA-MB-231 cells treated with a vehicle or the FAKi VS-4718 (1 μM) for 24 hours followed by Western blotting for TGFβRI and TGFβRII. **d,** IP of FLAG from the lysis of MDA-MB-231 cells treated with a vehicle or the FAKi VS-6063 (10 μM, 24 hours), followed by Western blotting for phospho-Tyrosine (p-Tyr). **e,** Mass spectrometric analysis showing integrin β1 tyrosine (Y)–182 phosphorylation. **f,** The orientation of Y182 was illustrated in the cartoon diagram of TBFβRI structure (PDB ID: 1ias) as generated using PyMOL. The GS domain of TBFβRI is colored in yellow. **g,** Close-up view of the interaction interfaces of docked complex model between TGFβRII (PDB ID: 5e92) and TGFβRI (PDB ID: 1ias) without (top)/with (bottom) Y182 phosphorylation, obtained through protein-protein docking using ClusPro server. **h,** HEK 293FT cells were transfected with TGFβRII, and FLAG tagged wild type TGFβRI (RI-WT-FLAG) or TGFβRI-Y182F mutant (RI-YF-FLAG). IP of FLAG from the lysis of cells using anti-FLAG antibody, followed by Western blotting for TGFβRII.

Structural analysis revealed that Y182 of TGFβRI is highly exposed for potential phosphorylation as described above (Fig. 4f), and it is close to two threonine and one serine sites (T185, T186, S187) in TGFβRI (Fig. 4g), which are bound and phosphorylated by TGFβRII ^29,32^. Our protein docking suggests that Y182 of TGFβRI is oriented toward K381 of TGFβRII with a distance of 4.1 Å (Supplementary Fig. 4i left, and Fig. 4g top). Upon phosphorylation, the distance of the negatively charged phosphate group of TGFβRI to positively charged K381 of TGFβRII becomes much closer (2.1 Å), therefore, significantly enhancing binding of TGFβRI to TGFβRII through increased charge-charge interactions with positively charged K381, as well as Mg^2+^ coordinated with ATP (Supplementary Fig. 4i right and Fig. 4g bottom). Consequently, such increased TGFβRI binding to TGFβRII may further promote the phosphate transfer from the TGFβRII-bounded ATP to T185, T186 and S187 in the GS domain of TGFβRI (Fig. 4g). To determine whether phosphorylation of Y182 at TGFβRI changes the binding affinity of TGFβRI to TGFβRII, we stably expressed FLAG-tagged wild-type TGFβRI (RI-WT-FLAG), a non-phosphorylatable Y182F mutant TGFβRI (RI-YF-FLAG), or a phosphomimic Y182D mutant TGFβRI (RI-YD-FLAG) in HEK 293FT cells. The wild-type TGFβRI, mutant TGFβRI Y182F or TGFβRI Y182D were pulled down with an anti-FLAG antibody followed by Western blotting of TGFβRII in these cells after TGFβ treatment. We found that the non-phosphorylatable TGFβRI-Y182F mutant had reduced binding to TGFβRII, compared with wild-type TGFβRI and TGFβRI-Y182D mutant (Fig. 4h and Supplementary Fig. 4j). The data clearly indicated that the phosphorylation of TGFβRI at Y182 enhanced TGFβRI binding to TGFβRII in response to TGFβ stimulation, which is consistent to our protein docking analysis. In summary, EZH2-activated FAK can phosphorylate TGFβRI at Y182, which alters amino acid interaction patterns to enhance the binding to TGFβRII resulting in activation of TGFβRI.

### EZH2 increases the FAK upstream *ITGB1* expression

FAK signaling is initiated by integrin-mediated cell adhesions. Integrins (such as β1 or β3) can facilitate FAK autophosphorylation at tyrosine 397, which increases the catalytic activity of FAK^22,33^. To understand the mechanism underlying increased pY397-FAK by EZH2, we investigated whether and how EZH2 regulates expression of integrins β1 or β3. Since EZH2 methyltransferase activity is not required for increasing pY397-FAK (Fig. 3f) and we recently reported that EZH2 can function as a transcription co-factor of RNA Pol II to upregulate mRNA transcription^34^, we examined whether EZH2 also regulate RNA Pol II transcription of genes encoding β1, β3, or other genes that may regulate pY397-FAK. We performed IP RNA Pol II from EZH2-expressing MDA-MB-231 cells versus from 231.KO#1 cells and followed by chromatin IP sequencing (ChIP-seq). The data showed that knocking out EZH2 in 231.KO#1 cells decreased RNA Pol II bindings at the promoter regions of 470 genes to levels lower than those in EZH2-expressing MDA-MB-231 cells (Table 2)^34^. Of the 470 genes, *ITGB1* (encoding integrin β1) is substantially decreased (Fig. 5a), whereas *ITGB3* is not (encoding integrin β3) (Supplementary Fig. 5a). Consistently, qRT-PCR showed that knockdown and knockout of EZH2 downregulated *ITGB1* mRNA expression (Fig. 5b, c), resulting in decreased integrin β1 protein expression (Supplementary Fig. 5b, c), while knockdown of EZH2 did not affect β3 mRNA expression (Supplementary Fig. 5d). Also, *ITGB1*-promoter luciferase reporter assays showed that *ITGB1* promoter activity was higher in EZH2-expressing MDA-MB-231 and 231.sgCtrl cells than in EZH2-null cells (231.KO#1 and 231.KO#2) (Fig. 5d).

**Figure 5.**
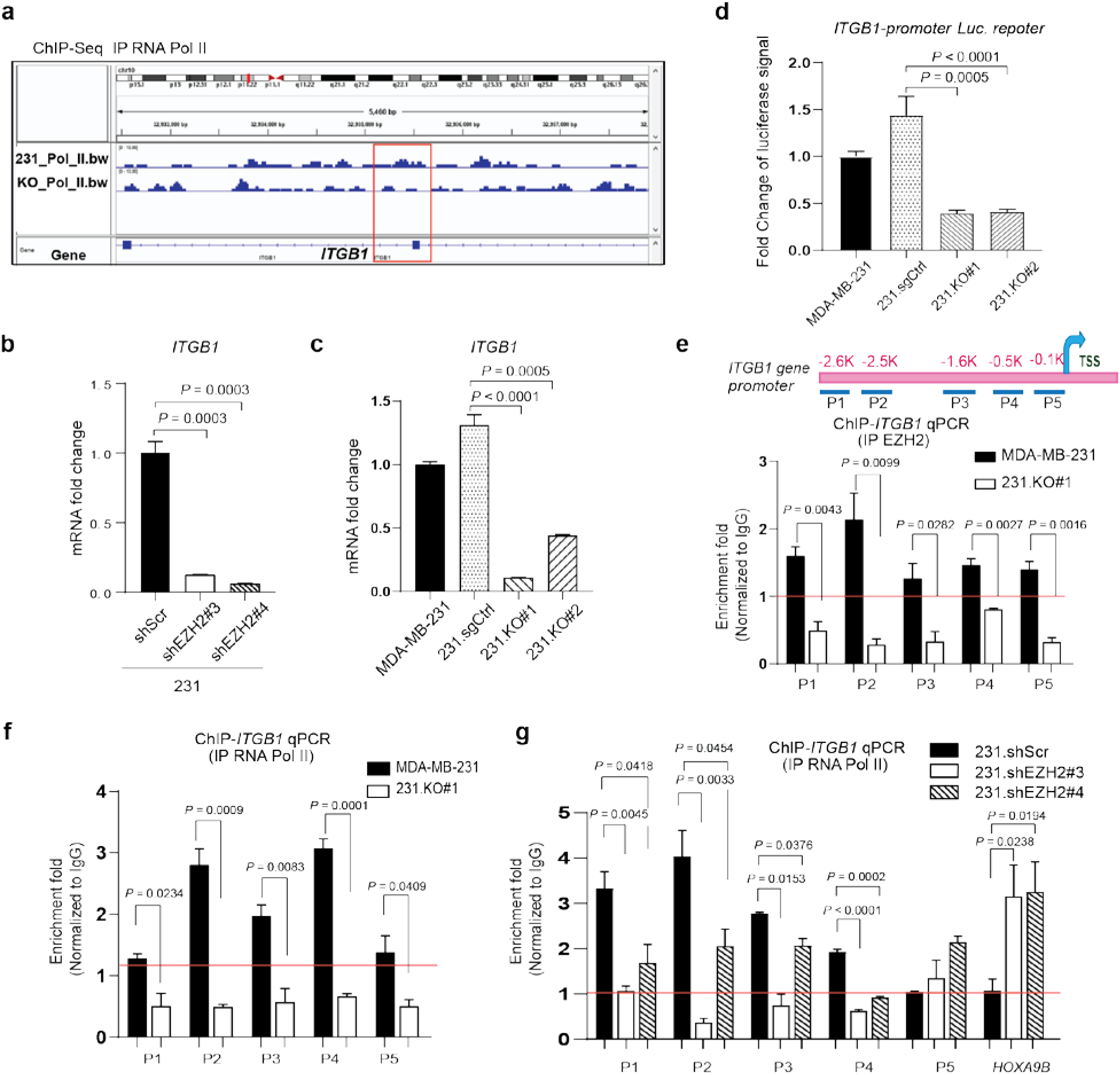
EZH2 increases the FAK upstream *ITGB1* expression. **a,** Screenshot of the RNA Pol II ChIP sequencing (ChIP-seq) signal at the *ITGB1* promoter locus in MDA-MB-231 (231_Pol_II.bw) and 231.KO#1 (KO_Pol_I.bw) cells. **b,** qRT-PCR analysis of *ITGB1* mRNA expression in the indicated cells. Data are presented as means ± S.E.M. (*t*-test). **c,** qRT-PCR analysis of *ITGB1* mRNA expression in the indicated cells. Data are presented as means ± S.E.M. (*t*-test). **d,***ITGB1* promoter activity in MDA-MB-231, 231.sgCtrl, 231.KO#1, and 231.KO#2 cells as measured using a dual-luciferase reporter assay. ITGB1 firefly luciferase signal was divided by the control renilla luciferase signal, and the ratios of luciferase/renilla in four cell lines were normalized to that of MDA-MB-231 cells. *n* ≥ 3. Data are means ± SEM, *t* test. **e,** Top: the locations of the primers at the *ITGB1* gene promoter area for ChIP-qPCR. TSS, transcription start site. Bottom: EZH2 was immunoprecipitated from MDA-MB-231 and 231.KO#1 cells, and EZH2 binding to *ITGB1* in the cells was detected using qPCR with the indicated primers. All fold-enrichment values were normalized according to IgG values. Data are presented as means ± S.E.M. (*t*-test). **f,** RNA Pol II was immunoprecipitated from MDA-MB-231 and 231.KO#1 cells, and RNA Pol II binding to *ITGB1* in the cells was detected using qPCR with the indicated primers. All fold-enrichment values were normalized according to IgG values. Data are presented as means ± S.E.M. (*t*-test). **g,** RNA Pol II was immunoprecipitated from 231.shScr, 231.shEZH2#3, and 231.shEZH2#4 cells, and RNA Pol II binding to *ITGB1* or *HOXA9B* in the cells was detected using qPCR with the indicated primers. *HOXA9B* was used as a negative control. All fold-enrichment values were normalized according to IgG values. Data are presented as means ± S.E.M. (*t*-test).

We further performed ChIP-qPCR to detect EZH2 binding and RNA Pol II binding to the *ITGB1* promoter in MDA-MB-231 versus 231.KO#1 cells using a series of PCR primers that bind to various regions of the *ITGB1* promoter from −2.6 kb upstream of (primer P1) to near (primer P5), the *ITGB1* transcription start site (Fig. 5e, top). In MDA-MB-231 cells, EZH2 was recruited to the *ITGB1* promoter from P1 to P5 loci, and expectedly, in 231.KO#1 cells, binding of EZH2 to the *ITGB1* promoter was lost (Fig. 5e, bottom). Notably, RNA Pol II bound well to P1 to P5 loci within the *ITGB1* promoter that overlapped with EZH2 binding loci in MDA-MB-231 cells, but RNA Pol II binding to the *ITGB1* promoter was also lost in 231.KO#1 cells without EZH2 (Fig. 5f), indicating EZH2 is required for RNA Pol II binding to the *ITGB1* promoter. These ChIP-qPCR data on RNA Pol II were consistent with ChIP-seq data (Fig. 5a). Similarly, shRNA-mediated knocking down of EZH2 in 231.shEZH2#3 and 231.shEZH2#4 cells also reduced the RNA Pol II binding at P1 to P4 loci of the *ITGB1* promoter compared to control 231.shScr cells (Fig. 5g), which paralleled with the reduced EZH2 binding (Supplementary Fig. 5e). *HOXA9B* is a well-known methyltransferase substrate of EZH2^35^. Expectedly, EZH2 knockdown in 231.shEZH2#3 and 231.shEZH2#4 cells reduced both EZH2 binding and H3K27me3 binding, but increased RNA Pol II binding at the *HOXA9B* promoter, compared to control 231.shScr cells; However, H3K27me3 binding to the *ITGB1* promoter were similar in control 231.shScr cells versus EZH2-knockdown cells (Supplementary Fig. 5f), further indicating EZH2 regulated *ITGB1* independent of its methyltransferase function. Taken together, both EZH2 and RNA Pol II bind to the same promoter regions of *ITGB1*, and EZH2 is likely functioning as a co-factor of RNA Pol II to upregulate *ITGB1* transcription independent of its methyltransferase function.

Since integrin β1 is responsible for FAK activation^22^ and our data confirmed that activated FAK bound to and phosphorylated TGFβRI (Fig. 4a, d and e), we questioned whether integrin β1 can bind to TGFβRI in the same complex. IP integrin β1 followed by Western blotting of TGFβRI and reverse IP TGFβRI followed by Western blotting of integrin β1 clearly showed that integrin β1 can bind to TGFβRI in MDA-MB-231 cells (Supplementary Fig. 5g, h), which further demonstrate the cross interactions between the TGFβ/TGFβRI pathway and the integrin β1/FAK pathway.

### Treatment with a clinically applicable FAK inhibitor blocks EZH2-induced breast cancer bone metastasis

Our finding that EZH2, via upregulating integrin β1 transcription, activated FAK that stimulated the TGFβ/Smad2 pathway to increase bone metastasis prompted us to test FAK inhibitor for treatment of bone metastases of high EZH2 expressing breast cancers. For bone metastasis outgrowth model, GFP- and luciferase-labeled MDA-MB-231 cells, which have relatively high EZH2 expression among tested breast cancer cell lines (Supplementary Fig. 6a), were intratibially injected into nude mice. We treated these mice with FAKi VS-6063 (50 mg/kg twice a day, oral gavage), which is currently tested in clinical trials for treating patients with advanced lymphoma or solid tumors (ClinicalTrials.gov Identifier: NCT04439331, NCT03875820). To confirm that EZH2-induced breast cancer bone metastasis outgrowth is independent of its methyltransferase function, a group of mice was treated with EZH2 methyltransferase inhibitor GSK126 (100 mg/kg once a day, i.p. injection). The resulting bone metastasis outgrowth were detected using BLI, which confirmed that GSK126-based treatment did not block tumor outgrowth in the bones (Fig. 6a). Excitingly, treatment with the FAKi VS-6063 impeded the outgrowth of bone tumors significantly more than in the control group (*P* = 0.0442) (Fig. 6A) and did not induce significant side effects (Supplementary Fig. 6b-e). IHC staining of bone metastasis showed that FAKi VS-6063, but not GSK126, significantly reduced pS465/467 Smad2 (*P* = 0.0066) level and PTHLH expression (*P* = 0.0074) in the bone metastases and both drugs effectively inhibited their targets (Fig. 6b, c and Supplementary Fig. 6f).

**Figure 6.**
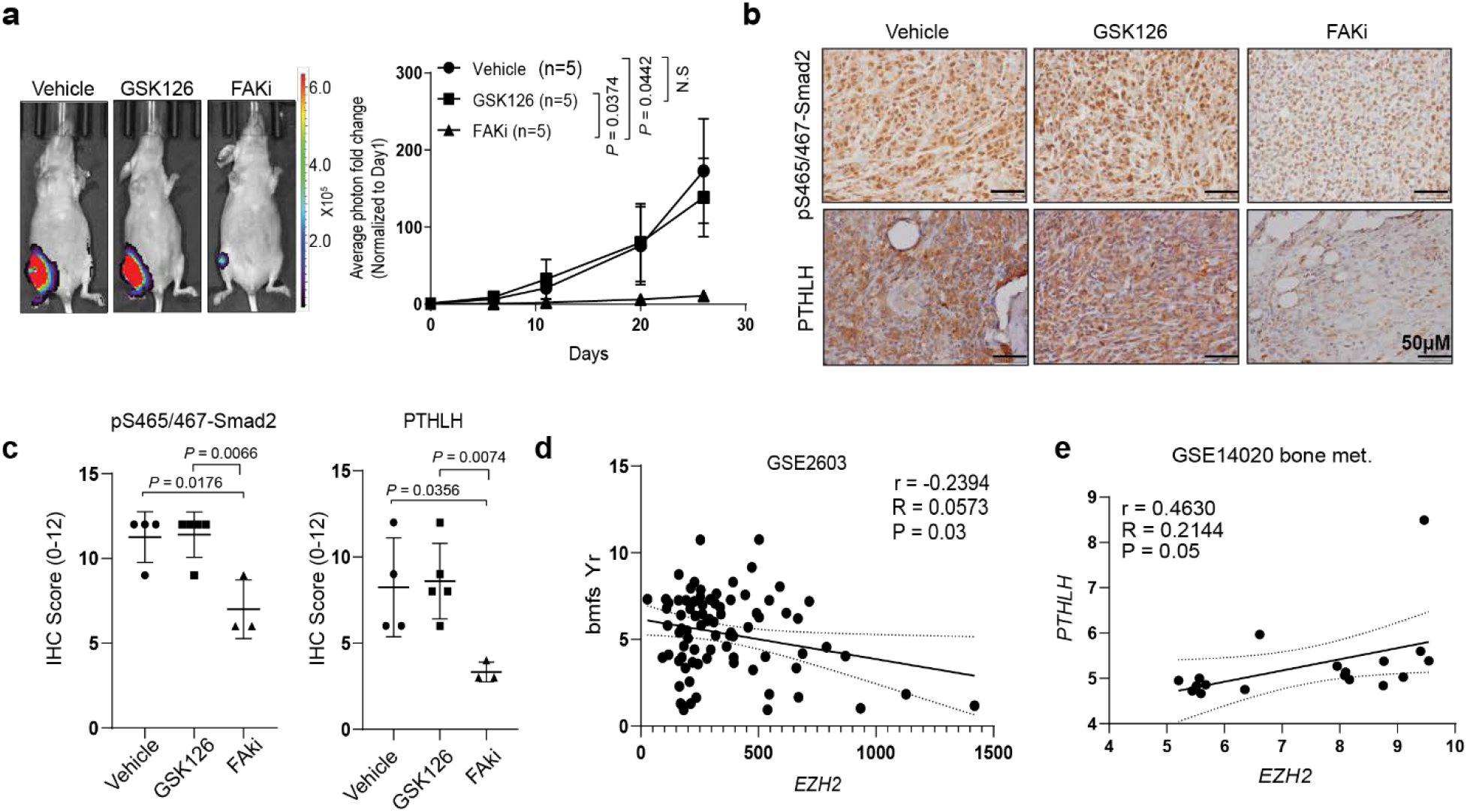
Treatment with a clinically applicable FAK inhibitor blocks EZH2-induced breast cancer bone metastasis. **a,** Representative bioluminescence images of and quantification of the fold change in BLI intensity in the right leg region in three subgroups of mice intratibially injected with MDA-MB-231 cells and given treatment with a vehicle, GSK126 (100 mg/kg/day, i.p. injection), or the FAKi VS-6063 (50 mg/kg, twice a day, oral gavage) beginning on day 18 of injection. The BLI signal was normalized according to the signal on the first day after injection. Data are presented as means ± S.D. (*t*-test). **b and c,** Representative pictures of IHC staining and quantification of pS465/467-Smad2 and PTHLP expression in the bone metastasis samples obtained from the three subgroups of mice in **a**. **d,** The Pearson *r* correlation for *EZH2* RNA expression and bone metastasis-free survival years (bmfs Yr) in breast cancer patients (GSE2603 data set). **e,** The Pearson *r* correlation for *EZH2* and *PTHLH* RNA mRNA expression in bone metastasis samples obtained from breast cancer patients (GSE14020 data set).

Finally, we examined the GSE2603 data set and validated that EZH2 expression was negatively correlated with bone metastasis-free survival in breast cancer patients (*r* = −0.2394, *P* = 0.03) (Fig. 6d), suggesting that high EZH2 expression in primary breast tumors produces a high risk of developing bone metastasis in patients. We also examined the correlation between the expression of *EZH2* and the downstream effector *PTHLH* in bone metastasis tissues obtained from breast cancer patients in the GSE14020 data set. We found that *EZH2* mRNA expression was positively correlated with *PTHLH* mRNA expression in bone metastases (*r* = 0.4630, *P* = 0.05) (Fig. 6e) but not in metastases to other organ sites (e.g., lung, liver, brain metastases; *r* = 0.2452, *P* = 0.097) (Supplementary Fig. 6g). These clinical data confirmed that EZH2 high expression can increase *PTHLH* expression that promotes bone metastasis in patients.

## Discussion

As described herein, we revealed a mechanism of how EZH2 promotes breast cancer bone metastasis. Specifically, EZH2 works as a transcription co-factor of RNA Pol II to increase *ITGB1* gene transcription; the increased integrin β1 induces phosphorylation of Y397 on FAK leading to FAK activation; activated pY397-FAK phosphorylates TGFβRI at Y182 that increases TGFβRI’s binding affinity for TGFβRII in response to TGFβ exposure, thereby triggering pS465/467-Smad2 that induces the downstream effector PTHLH; PTHLH accelerates osteolysis and thus driving the feed-forward vicious cycle of breast cancer bone metastasis outgrowth (Fig. 7). Since FAK and TGFβ enhances epithelial-mesenchymal transition and cell migration^8,36^, EZH2-induced FAK/TGFβ signaling activation is also an underlying mechanism of the strong migration and invasion ability of EZH2 high-expressing breast cancer cells.

**Figure 7.**
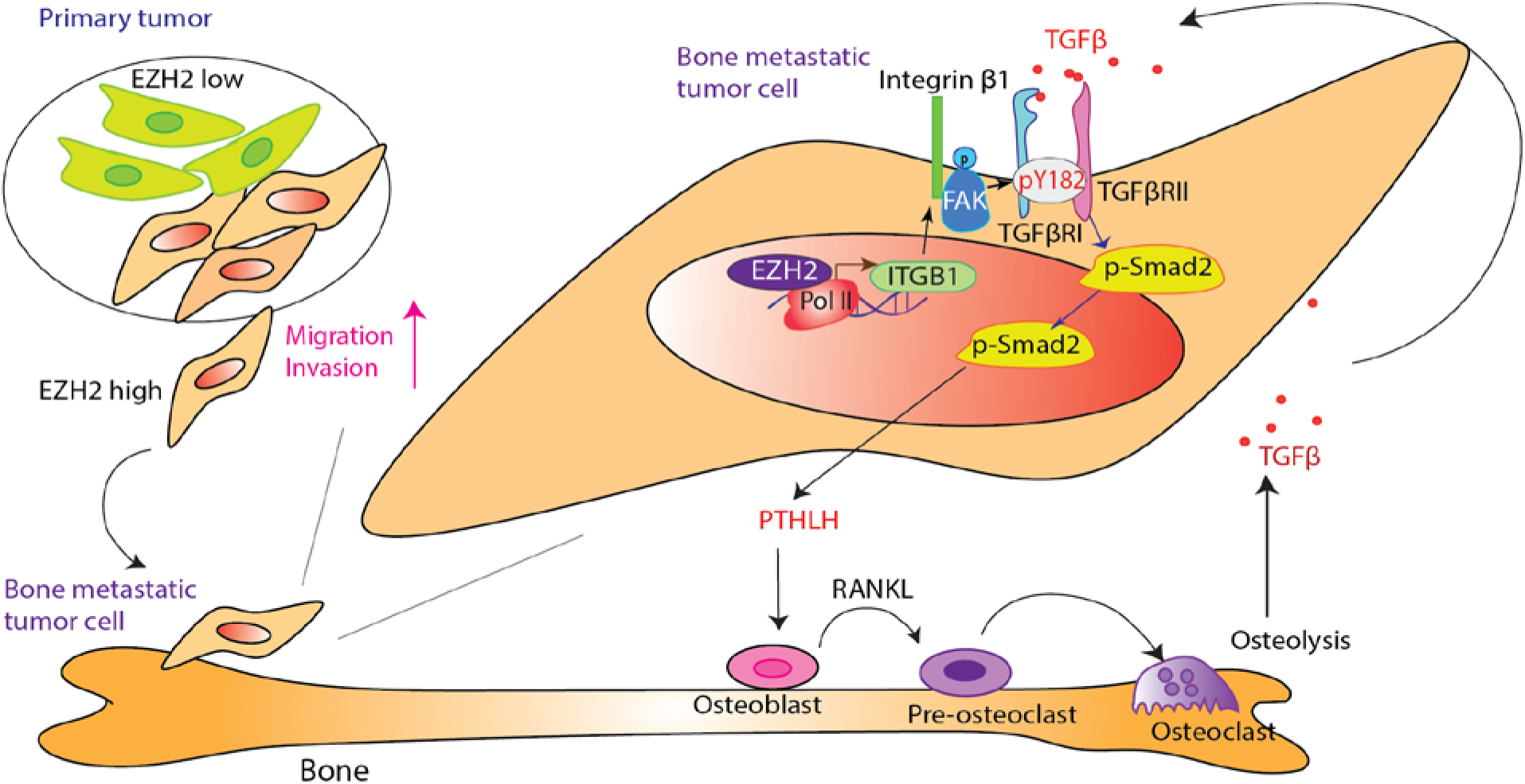
A model of EZH2’s interaction with TGFβ signaling in enhancing breast cancer bone metastasis. High EZH2-expressing cells have strong ability to invade from the primary tumor to the bone. In bone-metastatic tumor cells, EZH2 works as a transcription co-factor to increase ITGB1 transcription. Integrin β1 activates FAK at Y375, which phosphorylates TGFβRI at Y182. pY182-TGFβRI increases the binding of TGFβRI to TGFβRII, thereby activating pS465/467-Smad2, PTHLH expression, and the entire vicious cycle of breast cancer bone metastasis.

Although the function of TGFβ signaling in the bone metastasis of breast cancer is well known, the cross-talk between the integrin/FAK and TGFβ pathways is not well documented. It was reported that TGFβ activated FAK through integrin β3 or β1 and leading to p38 MAPK activation in renal cell carcinoma and hepato-carcinoma cells^37,38^. In the present study, we found that integrin β1-FAK is involved in the classical TGFβ/Smad-dependent pathway rather than the p38 MAPK pathway. Also, our data on the binding between integrin β1 and TGFβRI suggested that TGFβ may activate FAK through the TGFβRI- integrin β1 complex. Administering FAKi and genetically rendering FAK deficient in breast cancer cells abrogated the interaction between TGFβRI and TGFβRII and thereby blocked phosphorylation of Smad2 and expression of its downstream effector PTHLH. Our study demonstrated that the integrin/FAK and TGFβ pathways can cross talk and critically cooperate in driving the feed-forward vicious cycle of breast cancer bone metastasis.

It was reported that FAK and integrin β3 bind to TGFβRII in stellate hepatic cells^39^ and breast cancer cells^40^. Here, we detected that in EZH2 high expressing breast cancer cells, FAK and integrin β1 bound to TGFβRI rather than TGFβRII and that FAK phosphorylated TGFβRI. Little is known about tyrosine phosphorylation sites in TGFβRI and their functions, although several serine phosphorylation sites have been reported in TGFβRI^41^. Our mass spectrometry analysis identified an unreported tyrosine phosphorylation site at Y182 that fits the SH2 motif, and protein structure analysis and IP/Western blotting experiments showed that the Y182 of TGFβRI is important to regulating the binding of TGFβRI to TGFβRII and subsequent TGFβ/Smad2 pathway activation. Our findings revealed a new regulation mechanism of TGFβ/Smad2 pathway activation signifying the intricate regulations of TGFβ pathway.

EZH2 is a classic epigenetic protein that silences tumor suppressors through H3K27me3^13^. Recently, its noncanonical roles in the development of various cancers are gaining increasing attention. For example, EZH2 can methylate non-histone substrates, such as Jarid2, STAT3, RORα, and PLZF, to regulate their transcription function or protein stability^42–45^. EZH2 also have functions independent of its histone methyltransferase activities. For example, EZH2 forms a complex with RelA and RelB to activate nuclear factor κB signaling in estrogen receptor-negative breast cancer cells^46^ and activates androgen receptor gene transcription through binding at the androgen receptor promoter in prostate cancer cells^47,48^. We recently reported that EZH2 can function as a methyltransferase-independent transcription factor to upregulate *c-JUN* expression that induced G-CSF to facilitate the brain infiltration of immunosuppressive neutrophils^34^. In the present study, we found that EZH2 upregulated *ITGB1* transcription in breast cancer cells by functioning as a transcriptional co-factor of RNA Pol II to facilitate its binding to the *ITGB1* promoter. Thus, *ITGB1* is a newly identified substrate regulated by EZH2 methyltransferase-independent activity.

EZH2 is highly expressed in various human malignancies and regulates tumor progression. Therefore, it is regarded as an attractive therapeutic target in cancer patients^49^. However, targeting EZH2 with methyltransferase inhibitors has not always proven to be beneficial in clinical trials^49–52^, partially because of the EZH2 methyltransferase-independent functions in promoting cancer development as mentioned above. In the present study, we found that the small-molecule EZH2 inhibitor cannot block MDA-MB-231 cell-induced bone metastasis. However, targeting EZH2 downstream effector FAK with clinically applicable kinase inhibitors have striking effects on blocking breast cancer bone metastasis. Since EZH2 plays distinct functions in different types of cancer and in metastases of different organs, targeting downstream effectors of EZH2, but not EZH2’s enzyme function, is an effective alternative strategy for treating high EZH2-expressing bone metastases.

In summary, our work showed that FAK is a downstream effector of EZH2 and is involved in the vicious cycle of breast cancer bone metastasis. Treatment with a FAKi combined with standard antiresorptive agents, chemotherapy, or radiotherapy may provide added benefit to breast cancer patients who suffer from bone metastasis.

## Methods

### Reagents and plasmids

Antibodies against EZH2 (#5246), H3 (#4499), pS465/467-Smad2 (#3108), Smad2 (#5339), Smad3 (#9513), Smad4 (#46535), pT180/Y182-p38 (#4511), p38 (#8690), pY397-FAK (#3283), FAK (#3285), and FLAG (#14793) were purchased from Cell Signaling Technology. Antibodies against β-actin (A5441) and α-tubulin (T6074) were purchased from Sigma-Aldrich. Antibodies against TGFβRI (ab31013) and TGFβRII (ab184948) were purchased from Abcam. An antibody against TGFβRI (#ABF17-I) was purchased from Millipore. Antibodies against integrin β1 (sc-8978) and IgG (sc-2025, sc-2027) were purchased from Santa Cruz Biotechnology. Antibodies against RNA Pol II (NB200-598) were purchased from Novus Biologicals. An antibody against phosphor-tyrosine (#610000) was purchased form BD Biosciences. An antibody against PTHLH (#MAB6734) was purchased from R&D Systems. GSK126 (#15415), VS-6063 (#17737) and VS-4718 (#17668) were purchased from Cayman Chemical. GSK126 (#HY-13470) and VS-6063 (#HY-12289) were purchased from MedChemExpress. TGFβ (#4342-5) was purchased from BioVison. A leukocyte acid phosphatase kit (#387A) for tartrate-resistant acidic phosphatase (TRAP) staining was purchased from Sigma-Aldrich. LipoD293 DNA *in vitro* transfection reagent (#SL100468) and pepMute small interfering RNA (siRNA) transfection reagent (#SL100566) were purchased from SignaGen.

A pLenti-HA-EZH2 lentiviral vector was provided by Dr. Mien-Chie Hung^34^. Lentiviral-based pLKO.1 short hairpin RNAs (shRNAs) targeting EZH2 (shEZH2#3 [TRCN0000040076] and shEZH2#4 [TRCN0000010475]) and FAK (shFAK#2 [TRCN0000121211], shFAK#3 [TRCN0000121318], and shFAK#21 [TRCN0000121321]) were purchased from Sigma-Aldrich. pRK5-TGFβRI-FLAG plasmid (#14833) and pCMV5B- TGFβRII plasmid (#11766) were purchased from Addgene.

### Cell lines and cell culture

The human breast cancer cell lines MDA-MB-231 and MCF7 and mouse mammary tumor cell line 4T1 were purchased from the ATCC. The MDA-MB-231 subline 231-1566 was provided by Dr. Hung’s lab^7,53^. HEK 293FT cells were purchased from Thermo Fisher Scientific. These cell lines were further characterized by The University of Texas MD Anderson Cancer Center Cytogenetics and Cell Authentication Core and were tested for mycoplasma contamination. Cancer cells were cultured in Dulbecco’s modified Eagle’s medium/F12 medium supplemented with 10% fetal bovine serum (FBS; Thermo Fisher Scientific; #SH3007103). The murine osteoblast cell line MC3T3 was obtained from Dr. Sue-Hwa Lin’s lab^7^ and maintained in α-minimum essential medium with 10% FBS. The murine preosteoclast cell line RAW 264.7, obtained from the ATCC, was maintained in Dulbecco’s modified Eagle’s medium/high-glucose medium with 10% FBS for regular culture.

### RNA interference, transient transfection, CRISPR/CAS9 knockout cell line generation

Transient transfection of siRNAs into cancer cells was performed using pepMute siRNA Transfection Reagent (SignaGen; #SL100566). SiRNAs targeting FAK (siFAK#1 [SASI_Hs01_00035697] and siFAK#2 [SASI_Hs01_00035698]) and proline-rich tyrosine kinase 2 (PYK2; siPYK2#44 [SASI_Hs01_00207544], siPYK#49 [2SASI_Hs01_00032249], and siPYK2#50 [SASI_Hs01_00032250]) were purchased from Sigma-Aldrich. In transient transfection of HEK293FT cells with pRK5-TGFβRI-FLAG plasmid (#14833) or pCMV5B-TGFβRII plasmid (#11766) using Lipofectamine 3000 transfection kit (Invitrogen, #L3000-008). For lentivirus production, shRNA gene knockdown or gene overexpression lentiviral vectors were transfected into HEK 293FT cells together with a packaging plasmid (psPAX2; Addgene; #12260) and envelope plasmid (pMD2G; Addgene; #12259) using LipoD293 reagent (SignaGen; SL100668) according to the manufacturer’s instructions. The lentiviruses were collected, filtered, and used to infect target cells in the presence of 8-10 μg/mL hexadimethrine bromide (Polybrene) for 24 hours. The infected cells were selected using 300 μg/mL hygromycin (InvivoGen; #ant-hg-1) for 10 days or 2 μg/mL puromycin (InvivoGen; #ant-pr-1) for 4 days to generate stable cell lines.

Gene-knockout cell lines were established as described previously^34^. Briefly, to generate EZH2-knockout (KO) MDA-MB-231 and 4T1 cells, pSpCAS9 (BB)-2A-Puro (PX459) V2 plasmids (Addgene; #62988) were used following the protocol described by Ran *et al*.^54^. A single guide RNA-targeting EZH2 was designed using the online CRISPR Design Tool (http://tools.genome-engineering.org) with the following primers: F5’- CACCGTGGTGGATGCAACCCGCAA-3’ and R5’-AAACTTGCGGGTTGCA TCCACCAC-3’. Single guide RNA-targeting EZH2 oligos underwent annealing and were inserted into a pSpCAs9 (BB)-2A-Puro (PX459) V2 plasmid followed by transformation of the plasmid into a Stbl3-competent *Escherichia coli* strain. The plasmids extracted from *E. coli* colonies were sequence-verified and transfected into cancer cells using Lipofectamine 2000 (Life Technologies; #11668030). After puromycin selection, single cells were expanded into subclones. EZH2 protein expression in the cells was detected using Western blotting, and EZH2 DNA modifications were validated via DNA sequencing.

### Site-specific mutation

Site-specific mutation was performed using a Q5 Site-Directed Mutagenesis Kit (New England BioLabs; #E0554S). EZH2 H689A mutation was using the primers F5’-CAAATGCTTCGGTAAATCCAAACTGC-3’ and R5’-CCGAAGCATTTGCAAAACGAATTTTG-3’. ITGB1 Y182F mutation was using the primers F5’- AGATTTAATTTTTGATATGACAACATCAGGG-3’ and R5’-TTTAAGGTGGTGCCCTCT-3’. ITGB1 Y182D mutation was using the primers AGATTTAATTGATGATATGACAACATCAGG-3’ and R5’- TTTAAGGTGGTGCCCTCT-3’.

### Western blotting and immunoprecipitation

Western blotting and immunoprecipitation (IP) were performed as described previously^55^. Briefly, for Western blotting, cells were lysed in lysis buffer (5 M urea, 10% sodium dodecyl sulfate [SDS], DNase-free water: 1:1:1) and then sonicated. The lysates were collected for Western blotting analysis. Proteins were separated using SDS-polyacrylamide gel electrophoresis and transferred onto a polyvinylidene difluoride membrane. After each membrane was blocked with 5% milk for 1 hour, it was probed with various primary antibodies overnight at 4°C followed by incubation with secondary antibodies for 1 hour at room temperature before being visualized with enhanced chemiluminescence reagent. For IP, cells were washed twice with phosphate-buffered saline and scraped into IP lysis buffer (1% Triton X-100, 150 mM NaCl, 10 mM Tris, pH 7.4, 1 mM EGTA, 1 mM EDTA, 0.5 mM sodium orthovanadate, 0.4 mM phenylmethanesulfonyl fluoride, 0.5% NP-40). The total cell lysates were precleared via incubation with protein G-linked agarose beads (Sigma; #1124323300) for 2 hours at 4°C. After preclearing, lysates were incubated with the primary antibody overnight at 4°C and then with protein G-linked agarose beads for 1 hour at 4°C. The beads were washed with IP buffer three times, and the protein immunocomplex was extracted from agarose and detected using SDS-polyacrylamide gel electrophoresis and Western blotting.

### Cell proliferation assays

Cell proliferation was measured using a 3-(4,5-dimethylthiazol-2-yl)-2,5-diphenyltetrazolium bromide (MTT) assay. Ten thousand MDA-MB-231 cells or 1000 4T1 cells per well (four wells per sample) were seeded in a 24-well plate, and the cell growth was examined via staining with MTT (Thermo Fisher Scientific; #M6496). The resulting intracellular purple formazan was solubilized using dimethyl sulfoxide and measured its absorbance on a plate reader at 570, measure also at 650 nm as reference wavelength, using a Gen5 microplate reader (BioTek); calculate the signal sample as OD570 minus OD650.

### Quantitative reverse transcription-polymerase chain reaction

Total RNA in cells was isolated using TRIzol reagent (Thermo Fisher Scientific; #15596026) and then reverse-transcribed using an iScript cDNA Synthesis Kit (Bio-Rad; #1708891). Quantitative reverse transcription (qRT)-polymerase chain reaction (PCR) analysis of cDNA expressions was conducted using SYBR FAST Universal qPCR Master Mix (Kapa Biosystems; #KK4602) with a StepOnePlus real-time PCR system (Applied Biosystems). Relative mRNA expression was quantified using the 2^-ΔΔCt^ method with logarithmic transformation. SYBR primers were obtained from Sigma or Integrated DNA Technologies (IDT). The following primers were used: human *PTHLH* (encoding PTHrP): F5’-TTTACGGCGACGATTCTTCC-3’, R5’-TTCTTCCCAGGTGTCTTGAG-3’; mouse *Pthlh* (encoding PTHLH): F5’- CATCAGCTACTGCATGACAAGG-3’, R5’-GGTGGTTTTTGGTGTTGGGAG-3’; human *interleukin-8* (*IL-8*; *CXCL8*): F5’-GAGTGATTGAGAGTGGACCACACT-3’, R5’- AGACAGAGCTCTCTTCCATCAGAAA-3’; mouse *Il-8* (*Cxcl15*): F5’- TCCTGCTGGCTGTCCTTAAC-3’, R5’-ACTGCTATCACTTCCTTTCTGTTG-3’; human *ACTB*: F5’-CATGTACGTTGCTATCCAGGC-3’, R5’-CTCCTAATGTCACGCACGAT-3’; mouse *Actb*: F5’-TCCTCCTGAGCGCAAGTACTCT-3’, R5’-CGGACTCATCGTACTCCTGCTT-3’. Human *ITGB1* (encoding integrin β1) primer #1 (H_ITGB1_2) and *ITGB3* (encoding integrin β3) primer #1 (H_ITGB3_1) were purchased from Sigma-Aldrich.

### Chromatin IP-quantitative PCR

Chromatin IP (ChIP)-quantitative PCR (qPCR) was performed as described previously^34^. Briefly, cells were fixed with 37% formaldehyde (final, 0.5%), treated with glycine (final, 125 mM), washed, resuspended in ChIP lysis buffer (50 mM Tris, pH 8.1, 10 mM EDTA, 1% SDS, 1% protease inhibitor cocktail, 10 mM phenylmethanesulfonyl fluoride), and sonicated. Lysates containing soluble chromatin were incubated with antibodies against EZH2, RNA Pol II, H3K27me3, or IgG overnight at 4°C and then incubated for an additional 2 hours at 4°C with added protein G-linked agarose beads. The agarose bead-bound complexes were then washed, and the protein-chromatin complexes were extracted from the agarose beads with elution buffer. Reversal of the cross-linking of protein and DNA was performed by incubating the elution buffer with 10 mg/mL RNase A and 5 M NaCl overnight at 65°C followed by incubation with 0.5 M EDTA, 1 M Tris, pH 6.5, and 10 mg/mL proteinase K for 1 hour at 45°C. Co-precipitated DNA was purified using a QIAquick spin column (QIAGEN), and 2 μL of DNA was analyzed via qPCR using specific primers for the *ITGB1* promoter. The ChIP assay primers used were as follows: ChIP-*ITGB1*_1F: 5’- GCAAGCTCAGGCATAACAGC-3’; ChIP-*ITGB1*_1R: 5’- CCCTGGCTCAGAGAGAATGC-3’; ChIP-*ITGB1*_2F: 5’-AGCCCTTGAAGATGGAGGTCT- 3’; ChIP-*ITGB1*_2R: 5’-AGACAATGAGGGCCATTTGTTTTT-3’; ChIP-*ITGB1*_3F: 5’- TTCTCGCAGCCATCTGCTAT-3’; ChIP-*ITGB1*_3R: 5’-GCCACTGGTTGCTGACTTGA-3’; ChIP-*ITGB1_*4F: 5’-CTGGATATGCTGGTCTGGGC-3’; ChIP-*IGB1*_4R: 5’- CCCAGAATCCATTCGTGCCT-3’; ChIP-*ITGB1*_5F: 5’-TGCGCTTTGACCAGTTAGGT-3’; ChIP-*ITGB1*_5R: 5’-GGAGCCTGACCATGAAGGAA-3’; *HOXA9B_*F: 5’- TCGCCAACCAAACACAACAGTC-3’; and *HOXA9B_*R: 5’-AAAGGGATCGTGCCGCTCTAC-3’. All fold enrichment values were normalized according to those of IgG. *HOXA9B* was used as a positive control for EZH2 and H3K27me3 binding.

### Triple co-culture assay and TRAP staining

Triple co-culture assay and TRAP staining were performed as described previously^7^. Murine RAW 264.7 preosteoclasts (3 × 10^4^ cells/well) were seeded directly into the wells of six-well co-culture plates, and MC3T3 cells (3 × 10^4^ cells/well) were seeded into Millicell Hanging Cell Culture Inserts (Millipore) in the six-well co-culture plates. The next day, MC3T3 cells attached to the membrane**s** of the inserts, and luciferase/green fluorescent protein (GFP)-labeled (GFP^+^) MDA-MB-231, 231.sgCtrl, 231.KO#1, or 231.KO#2 cells (3 × 10^4^ cells/well each) or 4T1, 4T1.KO#1, or 4T1.KO#2 cells (500 cells/well each) were added on top of the MC3T3 cell layer in triplicate and treated with 5 ng/mL TGFβ, 2 μM GSK126, or a vehicle. Co-culture assays were performed in Dulbecco’s modified Eagle’s medium/high-glucose medium supplemented with 10% FBS and that was changed every 2 days. TRAP staining of osteoclasts was performed on day 6 using a leukocyte acid phosphatase kit (Sigma-Aldrich; #387A). TRAP^+^ multinucleated cells were scored as mature osteoclasts and quantified. MC3T3 cells and GFP^+^ tumor cells were trypsinized from the inserts and calculated GFP^+^ cell numbers using flow cytometry.

### Luciferase reporter assay

Luciferase reporter assay was performed as described previously^34^. pGL4.10 (Luc.2; E665A) was purchased from Promega. The pGL4.10-*ITGB1* reporter and a control Renilla luciferase vector were co-transfected into breast cancer cell lines using Lipofectamine 3000 transfection kit (Invitrogen; #L3000-008). After 48 hours, luciferase activity was measured using a Dual-Luciferase Reporter Assay kit (Promega; E1910) with a 20/20 Luminometer (Turner Biosystems). The *ITGB1* promoter was generated via amplification of a genomic DNA sequence with PCR using the designed primers and then inserted upstream of the luciferase reporter gene in the pGL4.10 vector. The primer sequences for the *ITGB1*_reporter_1 were F5’-CGGGGTACCCTGGCTAATTTTTAGTAGAG-3’ and R5’-CCGGATATC ACCTAACTGGTCAAAGCGCA-3’.

### Flow cytometry

For detecting TGFβRI on cell surfaces, breast cancer cells were seeded at the same density and collected when they reached 80-90% confluence. One million cells in each sample were washed twice in fluorescence-activated cell sorter (FACS) buffer (1% bovine serum albumin in phosphate-buffered saline), resuspended in 100 μL of FACS buffer, and stained with 1 μg of an anti-TGFβRI antibody or mouse anti-IgG antibody for 1 hour, washed twice in FACS buffer, and stained with 1 μg of an APC anti-mouse secondary antibody for 1 hour. Afterward, cells were washed with cold phosphate-buffered saline twice and analyzed using a FACSCanto II flow cytometer (BD Biosciences). For a triple co-culture assay, MC3T3 cells and GFP^+^ tumor cells were trypsinized from the inserts in tubes and washed twice in cold FACS buffer, resuspended in 400 μL of FACS buffer, and analyzed using a FACSCanto II flow cytometer.

### Migration and invasion assays

For a migration assay, breast cancer cells (30,000 cells/well) in FBS-free medium were plated in uncoated Transwell inserts. For an invasion assay, Transwell inserts were coated with 14.3% Matrigel for 1 hour. Breast cancer cells (30,000 cells/well) in FBS-free medium were plated in coated Transwell inserts. A medium containing 10% FBS was added to the lower compartment of Transwell, as a chemical attractant. After culturing for 18 hours in a migration assay or 24-30 hours in an invasion assay, the migrated and invaded cells at the bottom of the Transwell inserts were stained with crystal violet and counted under a light microscope.

### Reverse-phase protein array

RPPA analysis of MDA-MB-231, 231.sgCtrl, 231.KO#1, and 231.KO#2 cells after treatment with a vehicle or 5 ng/mL TGFβ for 2 hours was performed at the MD Anderson Functional Proteomics Reverse Phase Protein Array (RPPA) Core. Briefly, cellular proteins were denatured using 1% SDS, serially diluted, and spotted on nitrocellulose-coated slides. Each slide was probed with a validated primary antibody plus a biotin-conjugated secondary antibody. The signal obtained was amplified using a Dako Cytomation-catalyzed system and visualized in a DAB colorimetric reaction. The slides were analyzed using customized Microvigene software (VigeneTech Inc.). Each dilution curve was fitted using a logistic model (“SuperCurve Fitting” developed at MD Anderson)^56^ and normalized according to median polish. Data clustering and heat map creation were performed using Morpheus software (one minus Pearson’s correlation).

### Mass spectrometry

Liquid chromatography/tandem-mass spectrometry was used to identify phosphorylation sites of TGFβRI. HEK 293FT cells were transfected with pRK5-TGFβRI-FLAG plasmid (Addgene, #14833) using Lipofectamine 3000 transfection kit; HEK 293FT cell lysates were immunoprecipitated with an anti-FLAG antibody. After protein gel electrophoresis, TGFβRI band was excised from the gels and subjected to tryptic digestion. An aliquot of the tryptic digest (in 2 % acetonitrile/0.1% formic acid in water) was analyzed by LC/MS/MS on an Orbitrap Fusion™ Tribrid™ mass spectrometer (Thermo Scientific™) interfaced with a Dionex UltiMate 3000 Binary RSLCnano System. Peptides were separated onto an Acclaim™ PepMap ™ C18 column (75 µm ID x 15 cm, 2 µm) at flow rate of 300 nl/min. Gradient conditions were: 3%-22% B for 40 min; 22%-35% B for 10min; 35%-90% B for 10 min; 90% B held for 10 min,(solvent A, 0.1 % formic acid in water; solvent B, 0.1% formic acid in acetonitrile). The peptides were analyzed using data-dependent acquisition method, Orbitrap Fusion was operated with measurement of FTMS1 at resolutions 120,000 FWHM, scan range 350-1500 m/z, AGC target 2E5, and maximum injection time of 50 ms; During a maximum 3 second cycle time, the ITMS2 spectra were collected at rapid scan rate mode, with HCD NCE 34, 1.6 m/z isolation window, AGC target 1E4, maximum injection time of 35 ms, and dynamic exclusion was employed for 20 seconds. The raw data files were processed using Thermo Scientific™ Proteome Discoverer™ software version 1.4, spectra were searched against the Uniprot-Homo sapiens database using the Mascot search engine v2.3.02 (Matrix Science) run on an in-house server. Search results were trimmed to a 1% FDR for strict and 5% for relaxed condition using Percolator. For the trypsin, up to two missed cleavages were allowed. MS tolerance was set 10 ppm; MS/MS tolerance 0.8 Da. Carbamidomethylation on cysteine residues was used as fixed modification; oxidation of methione as well as phosphorylation of serine, threonine and tyrosine was set as variable modifications.

### TGFβRI-TGFβRII complex modelling by molecular docking

High-resolution crystal structures of the cytoplasmic domain of TGFβRI (PDB ID: 1ias) and the kinase domain of TGFβRII (AMPPNP, an ATP analogue, bound state, PDB ID: 5e92) were obtained from Protein Data Bank. A phosphoryl group was added to TGFβRI Y182 using PyTMs^57^. The complex docking models between TGFβRI/pY182-TGFβRI and TGFβRII were built with ClusPro web-based server (http://cluspro.bu.edu/)^58^. The residues that were used as attraction restraint included T185, T186 and S187 of TGFβRI, and AMPPNP601 of TGFβRII was retained. The top 10 low-energy docked models from each restraint were downloaded from the server, and then visualized and analyzed with PyMOL. Two final models were selected on the basis of substrate recognition and phosphotransfer mechanism from ATP hydrolysis to TGFβRI.

### Animal experiments

All animal experiments were carried out in accordance with protocols approved by the MD Anderson Institutional Animal Care and Use Committee. Athymic NCr nu/nu mice were obtained from MD Anderson. The mice were exposed to a 12-hour light/12-hour dark cycle, bred as specific pathogen-free mice, and given free access to food and water. The number of mice used in each experimental group was determined via power analysis, and mice were grouped randomly for each experiment. All mice used were the same age (8 weeks) and had similar body weights. Two different injection models were used for bone metastasis studies. 1) Intracardiac injection model: 1 × 10^5^ cells of the 231-1566 sublines or EZH2-knockout MDA-MB-231 sublines were injected into the left ventricle in anesthetized female athymic NCr nu/nu mice. GSK126 was dissolved in 20% cyclodextrin (Captisol; CyDex Pharmaceuticals) and adjusted to a pH level of 4.0 to 4.5 with 1 N acetic acid following the instructions described by McCabe *et al*.^50^. GSK126 was administered to the mice via intraperitoneal (i.p.) injection three times a week at a dose of 150 mg/kg after 5 days of injections. 2) Intratibial injection model: 2 × 10^5^ MDA-MB-231 cells were injected into a tibia in anesthetized female athymic NCr nu/nu mice. VS-6063 was prepared in a vehicle (0.5% hydroxypropyl methylcellulose with 0.1% Tween 80) and administered to the mice via oral gavage (50 mg/kg) twice a day after 18 days of injection, and GSK126 was administered to them via i.p. injection every day at a dose of 100 mg/kg after 18 days of injections. Development of bone metastases was monitored using bioluminescence imaging (BLI). After anesthetized mice were intraperitoneally injected with 75 mg/kg D-luciferin, BLI was performed using a Xenogen IVIS 200 imaging system (PerkinElmer). Analysis of bone metastasis was performed using living image software by measuring the photon flux in the hindlimbs of mice. The photon flux data were normalized according to the signal on the day when mice were given the drug GSK126 or vehicle. Bone metastasis-free survival curves showed the time point at which each mouse experienced bone metastasis development according to threshold BLI signals in the hindlimbs. X-ray images of hindlimbs of mice were obtained using an IVIS Lumina XR system (PerkinElmer).

### Immunohistochemistry staining and scoring system

Standard immunohistochemistry (IHC) staining was performed as described previously^59^. The immunoreactive score (IRS) was used to quantify the IHC staining, ranging from 0 to 12 as a result of multiplication of positive cell proportion scores (0-4) and staining intensity scores (0-3). IHC staining and statistical analysis results were independently evaluated by two pathologists blinded to the experimental groups.

### Statistical analysis

All quantitative experiments were performed using at least three independent biological repeats, and the results are presented as means ± standard deviation (S.D.) or means ± standard error of the mean (S.E.M.). One-way analysis of variance (multiple groups) or *t*-tests (two groups) were used to compare the means for two or more samples using the Prism 8 software program (GraphPad Software). Survival was analyzed using Kaplan-Meier curves and log-rank tests. *P* values less than 0.05 (two-sided) were considered statistically significant.

## Acknowledgments

We thank members of the Yu’s laboratory for insightful discussions. We thank D. Norwood, the Department of Scientific Publications of MD Anderson Cancer Center for article revision. This work was supported by National Institutes of Health (NIH) grants R01CA184836 (D.Y.), R01 CA208213 (D.Y.), and R01CA231149 (D.Y.), the METAvivor grants 56675 and 58284 (D.Y.), China Scholarship Council 201706280072 (J.Q.) and NIH Cancer Center Support Grant P30CA016672 to MD Anderson Cancer Center (Functional Genomics Core, Flow Cytometry and Cellular Imaging resource, Advanced Technology Genomics Core, Research Histology Core Laboratory, Cytogenetics and Cell Authentication Core, Functional Proteomics Reverse Phase Protein Array (RPPA) Core, and Research Animal Support Facility-Houston). D. Yu is the Hubert L. & Olive Stringer Distinguished Chair in Basic Science at MD Anderson. Mass spectrum is performed at and supported in part by the Clinical and Translational Proteomics Service Center at the University of Texas Health Science Center.

## Authors’ Contributions

L.Z., J.Q. and D.Y. developed original hypothesis and designed experiments. L.Z., J.Q., Y.Q., Y.-W.H., Z.Z., P.L., J.Y., B.H., S.Z. and D.Y. performed experiments and/or analyzed data. L.Z., J. Q., and D.Y. wrote and edited the manuscript. D.Y. supervised the study.

**Correspondence and material requests** should be addressed to D.Y.

## References

1 Siegel, R. L., Miller, K. D. & Jemal, A. Cancer statistics, 2019. CA Cancer J Clin 69, 7–34, doi:10.3322/caac.21551 (2019).

2 Fornetti, J., Welm, A. L. & Stewart, S. A. Understanding the Bone in Cancer Metastasis. J Bone Miner Res 33, 2099–2113, doi:10.1002/jbmr.3618 (2018).

3 Rossi, L., Longhitano, C., Kola, F. & Del Grande, M. State of art and advances on the treatment of bone metastases from breast cancer: a concise review. Chin Clin Oncol 9, 18, doi:10.21037/cco.2020.01.07 (2020).

4 Mundy, G. R. Metastasis to bone: causes, consequences and therapeutic opportunities. Nat Rev Cancer 2, 584–593, doi:10.1038/nrc867 (2002).

5 Zhang, W., Bado, I., Wang, H., Lo, H. C. & Zhang, X. H. Bone Metastasis: Find Your Niche and Fit in. Trends Cancer 5, 95–110, doi:10.1016/j.trecan.2018.12.004 (2019).

6 Esposito, M., Guise, T. & Kang, Y. The Biology of Bone Metastasis. Cold Spring Harb Perspect Med 8, doi:10.1101/cshperspect.a031252 (2018).

7 Xu, J. et al. 14-3-3zeta turns TGF-beta’s function from tumor suppressor to metastasis promoter in breast cancer by contextual changes of Smad partners from p53 to Gli2. Cancer Cell 27, 177–192, doi:10.1016/j.ccell.2014.11.025 (2015).

8 Korpal, M. & Kang, Y. Targeting the transforming growth factor-beta signalling pathway in metastatic cancer. Eur J Cancer 46, 1232–1240, doi:10.1016/j.ejca.2010.02.040 (2010).

9 Kang, Y. Pro-metastasis function of TGFbeta mediated by the Smad pathway. J Cell Biochem 98, 1380–1390, doi:10.1002/jcb.20928 (2006).

10 Colak, S. & Ten Dijke, P. Targeting TGF-beta Signaling in Cancer. Trends Cancer 3, 56–71, doi:10.1016/j.trecan.2016.11.008 (2017).

11 Kang, Y. et al. Breast cancer bone metastasis mediated by the Smad tumor suppressor pathway. Proc Natl Acad Sci U S A 102, 13909–13914, doi:10.1073/pnas.0506517102 (2005).

12 Buijs, J. T., Stayrook, K. R. & Guise, T. A. The role of TGF-beta in bone metastasis: novel therapeutic perspectives. Bonekey Rep 1, 96, doi:10.1038/bonekey.2012.96 (2012).

13 Kim, K. H. & Roberts, C. W. Targeting EZH2 in cancer. Nat Med 22, 128–134, doi:10.1038/nm.4036 (2016).

14 Ren, G. et al. Polycomb protein EZH2 regulates tumor invasion via the transcriptional repression of the metastasis suppressor RKIP in breast and prostate cancer. Cancer Res 72, 3091–3104, doi:10.1158/0008-5472.CAN-11-3546 (2012).

15 Kowalski, P. J., Rubin, M. A. & Kleer, C. G. E-cadherin expression in primary carcinomas of the breast and its distant metastases. Breast Cancer Res 5, R217–222, doi:10.1186/bcr651 (2003).

16 Alford, S. H., Toy, K., Merajver, S. D. & Kleer, C. G. Increased risk for distant metastasis in patients with familial early-stage breast cancer and high EZH2 expression. Breast Cancer Res Treat 132, 429–437, doi:10.1007/s10549-011-1591-2 (2012).

17 Wang, J. et al. Alterations in enhancer of zeste homolog 2, matrix metalloproteinase-2 and tissue inhibitor of metalloproteinase-2 expression are associated with ex vivo and in vitro bone metastasis in renal cell carcinoma. Mol Med Rep 11, 3585–3592, doi:10.3892/mmr.2015.3164 (2015).

18 Quan, J., Hou, Y., Long, W., Ye, S. & Wang, Z. Characterization of different osteoclast phenotypes in the progression of bone invasion by oral squamous cell carcinoma. Oncol Rep 39, 1043–1051, doi:10.3892/or.2017.6166 (2018).

19 Soki, F. N., Park, S. I. & McCauley, L. K. The multifaceted actions of PTHrP in skeletal metastasis. Future Oncol 8, 803–817, doi:10.2217/fon.12.76 (2012).

20 Bendre, M. S. et al. Interleukin-8 stimulation of osteoclastogenesis and bone resorption is a mechanism for the increased osteolysis of metastatic bone disease. Bone 33, 28–37, doi:10.1016/s8756-3282(03)00086-3 (2003).

21 Kakonen, S. M. et al. Transforming growth factor-beta stimulates parathyroid hormone-related protein and osteolytic metastases via Smad and mitogen-activated protein kinase signaling pathways. J Biol Chem 277, 24571–24578, doi:10.1074/jbc.M202561200 (2002).

22 van Nimwegen, M. J. & van de Water, B. Focal adhesion kinase: a potential target in cancer therapy. Biochem Pharmacol 73, 597–609, doi:10.1016/j.bcp.2006.08.011 (2007).

23 Madan, R., Smolkin, M. B., Cocker, R., Fayyad, R. & Oktay, M. H. Focal adhesion proteins as markers of malignant transformation and prognostic indicators in breast carcinoma. Hum Pathol 37, 9–15, doi:10.1016/j.humpath.2005.09.024 (2006).

24 Lowe, C. et al. Osteopetrosis in Src-deficient mice is due to an autonomous defect of osteoclasts. Proc Natl Acad Sci U S A 90, 4485–4489, doi:10.1073/pnas.90.10.4485 (1993).

25 Soriano, P., Montgomery, C., Geske, R. & Bradley, A. Targeted disruption of the c-src proto-oncogene leads to osteopetrosis in mice. Cell 64, 693–702, doi:10.1016/0092-8674(91)90499-o (1991).

26 Zhang, X. H. et al. Latent bone metastasis in breast cancer tied to Src-dependent survival signals. Cancer Cell 16, 67–78, doi:10.1016/j.ccr.2009.05.017 (2009).

27 Nakao, A. et al. Identification of Smad7, a TGFbeta-inducible antagonist of TGF-beta signalling. Nature 389, 631–635, doi:10.1038/39369 (1997).

28 Heldin, C. H. & Moustakas, A. Signaling Receptors for TGF-beta Family Members. Cold Spring Harb Perspect Biol 8, doi:10.1101/cshperspect.a022053 (2016).

29 Wrana, J. L., Attisano, L., Wieser, R., Ventura, F. & Massague, J. Mechanism of activation of the TGF-beta receptor. Nature 370, 341–347, doi:10.1038/370341a0 (1994).

30 Lindfors, H. E., Drijfhout, J. W. & Ubbink, M. The Src SH2 domain interacts dynamically with the focal adhesion kinase binding site as demonstrated by paramagnetic NMR spectroscopy. IUBMB Life 64, 538–544, doi:10.1002/iub.1038 (2012).

31 Wu, J. C. et al. Focal adhesion kinase-dependent focal adhesion recruitment of SH2 domains directs SRC into focal adhesions to regulate cell adhesion and migration. Sci Rep 5, 18476, doi:10.1038/srep18476 (2015).

32 Wieser, R., Wrana, J. L. & Massague, J. GS domain mutations that constitutively activate T beta R-I, the downstream signaling component in the TGF-beta receptor complex. EMBO J 14, 2199–2208 (1995).

33 Naik, A. et al. Neuropilin-1 promotes the oncogenic Tenascin-C/integrin beta3 pathway and modulates chemoresistance in breast cancer cells. BMC Cancer 18, 533, doi:10.1186/s12885-018-4446-y (2018).

34 Zhang, L. et al. Blocking immunosuppressive neutrophils deters pY696-EZH2-driven brain metastases. Sci Transl Med 12, eaaz5387, doi:10.1126/scitranslmed.aaz5387 (2020).

35 Cao, R. & Zhang, Y. SUZ12 is required for both the histone methyltransferase activity and the silencing function of the EED-EZH2 complex. Mol Cell 15, 57–67, doi:10.1016/j.molcel.2004.06.020 (2004).

36 Sulzmaier, F. J., Jean, C. & Schlaepfer, D. D. FAK in cancer: mechanistic findings and clinical applications. Nat Rev Cancer 14, 598–610, doi:10.1038/nrc3792 (2014).

37 Feldkoren, B., Hutchinson, R., Rapoport, Y., Mahajan, A. & Margulis, V. Integrin signaling potentiates transforming growth factor-beta 1 (TGF-beta1) dependent down-regulation of E-Cadherin expression - Important implications for epithelial to mesenchymal transition (EMT) in renal cell carcinoma. Exp Cell Res 355, 57–66, doi:10.1016/j.yexcr.2017.03.051 (2017).

38 Cai, T., Lei, Q. Y., Wang, L. Y. & Zha, X. L. TGF-beta 1 modulated the expression of alpha 5 beta 1 integrin and integrin-mediated signaling in human hepatocarcinoma cells. Biochem Biophys Res Commun 274, 519–525, doi:10.1006/bbrc.2000.3177 (2000).

39 Chen, Y. et al. Focal Adhesion Kinase Promotes Hepatic Stellate Cell Activation by Regulating Plasma Membrane Localization of TGFbeta Receptor 2. Hepatol Commun 4, 268–283, doi:10.1002/hep4.1452 (2020).

40 Wendt, M. K. & Schiemann, W. P. Therapeutic targeting of the focal adhesion complex prevents oncogenic TGF-beta signaling and metastasis. Breast Cancer Res 11, R68, doi:10.1186/bcr2360 (2009).

41 Souchelnytskyi, S., ten Dijke, P., Miyazono, K. & Heldin, C. H. Phosphorylation of Ser165 in TGF-beta type I receptor modulates TGF-beta1-induced cellular responses. EMBO J 15, 6231–6240 (1996).

42 He, A. et al. PRC2 directly methylates GATA4 and represses its transcriptional activity. Genes Dev 26, 37–42, doi:10.1101/gad.173930.111 (2012).

43 Sanulli, S. et al. Jarid2 Methylation via the PRC2 Complex Regulates H3K27me3 Deposition during Cell Differentiation. Mol Cell 57, 769–783, doi:10.1016/j.molcel.2014.12.020 (2015).

44 Lee, J. M. et al. EZH2 generates a methyl degron that is recognized by the DCAF1/DDB1/CUL4 E3 ubiquitin ligase complex. Mol Cell 48, 572–586, doi:10.1016/j.molcel.2012.09.004 (2012).

45 Vasanthakumar, A. et al. A non-canonical function of Ezh2 preserves immune homeostasis. EMBO Rep 18, 619–631, doi:10.15252/embr.201643237 (2017).

46 Gonzalez, M. E. et al. Histone methyltransferase EZH2 induces Akt-dependent genomic instability and BRCA1 inhibition in breast cancer. Cancer Res 71, 2360–2370, doi:10.1158/0008-5472.CAN-10-1933 (2011).

47 Xu, K. et al. EZH2 oncogenic activity in castration-resistant prostate cancer cells is Polycomb-independent. Science 338, 1465–1469, doi:10.1126/science.1227604 (2012).

48 Kim, J. et al. Polycomb- and Methylation-Independent Roles of EZH2 as a Transcription Activator. Cell Rep 25, 2808–2820 e2804, doi:10.1016/j.celrep.2018.11.035 (2018).

49 Gulati, N., Beguelin, W. & Giulino-Roth, L. Enhancer of zeste homolog 2 (EZH2) inhibitors. Leuk Lymphoma 59, 1574–1585, doi:10.1080/10428194.2018.1430795 (2018).

50 McCabe, M. T. et al. EZH2 inhibition as a therapeutic strategy for lymphoma with EZH2-activating mutations. Nature 492, 108–112, doi:10.1038/nature11606 (2012).

51 Kondo, Y. Targeting histone methyltransferase EZH2 as cancer treatment. J Biochem 156, 249–257, doi:10.1093/jb/mvu054 (2014).

52 Italiano, A. et al. Tazemetostat, an EZH2 inhibitor, in relapsed or refractory B-cell non-Hodgkin lymphoma and advanced solid tumours: a first-in-human, open-label, phase 1 study. Lancet Oncol 19, 649–659, doi:10.1016/S1470-2045(18)30145-1 (2018).

53 Khotskaya, Y. B. et al. S6K1 promotes invasiveness of breast cancer cells in a model of metastasis of triple-negative breast cancer. Am J Transl Res 6, 361–376 (2014).

54 Ran, F. A. et al. Genome engineering using the CRISPR-Cas9 system. Nat Protoc 8, 2281–2308, doi:10.1038/nprot.2013.143 (2013).

55 Zhang, S. et al. SRC family kinases as novel therapeutic targets to treat breast cancer brain metastases. Cancer Res 73, 5764–5774, doi:10.1158/0008-5472.CAN-12-1803 (2013).

56 Hu, J. et al. Non-parametric quantification of protein lysate arrays. Bioinformatics 23, 1986–1994, doi:10.1093/bioinformatics/btm283 (2007).

57 Warnecke, A., Sandalova, T., Achour, A. & Harris, R. A. PyTMs: a useful PyMOL plugin for modeling common post-translational modifications. BMC Bioinformatics 15, 370, doi:10.1186/s12859-014-0370-6 (2014).

58 Kozakov, D. et al. The ClusPro web server for protein-protein docking. Nat Protoc 12, 255–278, doi:10.1038/nprot.2016.169 (2017).

59 Lu, J. et al. 14-3-3zeta Cooperates with ErbB2 to promote ductal carcinoma in situ progression to invasive breast cancer by inducing epithelial-mesenchymal transition. Cancer cell 16, 195–207, doi:10.1016/j.ccr.2009.08.010 (2009).

