## Supplemental figures for "EZH2 Engages TGFβ Signaling to Promote Breast Cancer Bone Metastasis via Integrin β1-FAK Activation"

### Supplementary Data

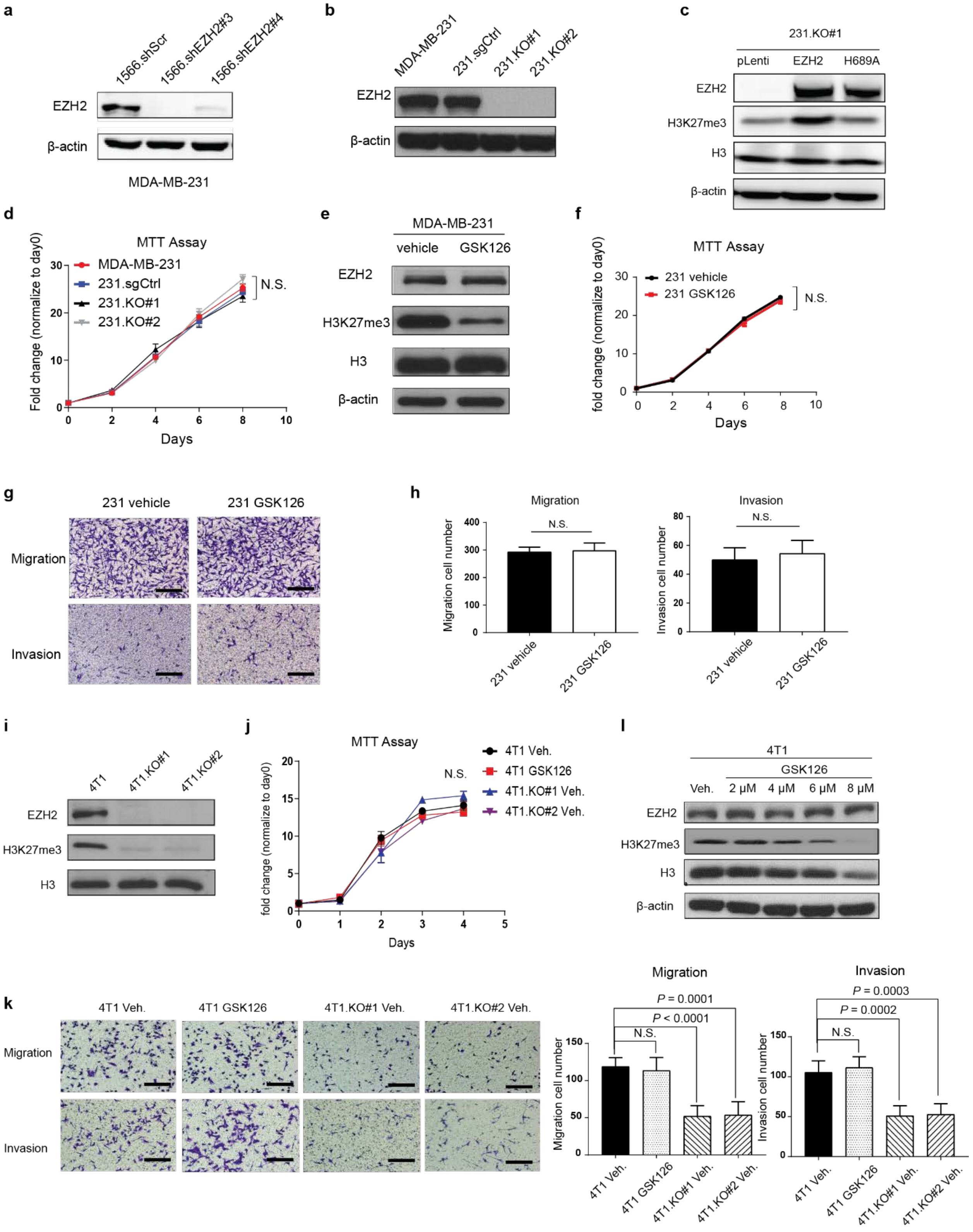

**Supplementary Fig. 1.** EZH2 promotes breast cance cell's migration and invasion. **a,** Western blotting of the expression of EZH2 and β-actin in 1566.shScr, 1566.shEZH2#3, and 1566.shEZH2#4 cells. **b,** Western blotting of the expression of EZH2 and β-actin in MDA-MB-231, 231.sgCtrl., 231.KO#1, and 231.KO#2 cells. **c,** Western blotting of the expression of EZH2, H3K27me3, H3, and β-actin in 231.KO#1 cell sublines with re-expression of a control pLenti vector (231.KO#1.pLenti), wild-type EZH2 (231.KO#1.EZH2), or EZH2 with the H689A mutant (231.KO#1.H689A). **d,** MTT assay results showing proliferation of MDA-MB-231, 231.sgCtrl., 231.KO#1, and 231.KO#2 cells. Data are presented as means ± S.D. (*t*-test). N.S., not significant. **e,** Western blotting of the expression of EZH2, H3K27me3, H3, and β-actin in MDA-MB-231 cells treated with a vehicle (DMSO) or GSK126 (2 μM, 24 hours). **f,** MTT assay results showing proliferation of MDA-MB-231 cells treated with a vehicle or GSK126 (2 μM, 24 hours). N.S., not significant. **g** and **h,** Representative images and quantification of invading and migrating MDA-MB-231 cells treated with a vehicle or GSK126 (2 μM, 24 hours). Scale bars, 100 μm. Data are presented as means ± S.D. (*t*-test). **i,** Western blotting of the expression of EZH2, H3K27me3, and H3 in 4T1, 4T1.KO#1, and 4T1.KO#2 cells. **j,** MTT assay results showing proliferation of 4T1 cells treated with vehicle or with GSK126 (6 μM), 4T1.KO#1 cells, and 4T1.KO2#2 cells treated with vehicle. N.S., not significant. **k,** Representative images and quantification of invading and migrating 4T1 cells treated with vehicle or GSK126 (6 μM), 4T1.KO#1 cells, and 4T1.KO#2 cells treated with vehicle. Scale bars, 100 μm. Data are presented as means ± S.D. (*t*-test). **l,** Western blotting of the expression of EZH2, H3K27me3, H3, and β-actin in 4T1 cells treated with a vehicle or GSK126 at different concentrations (2-8 μM, 48 hours).

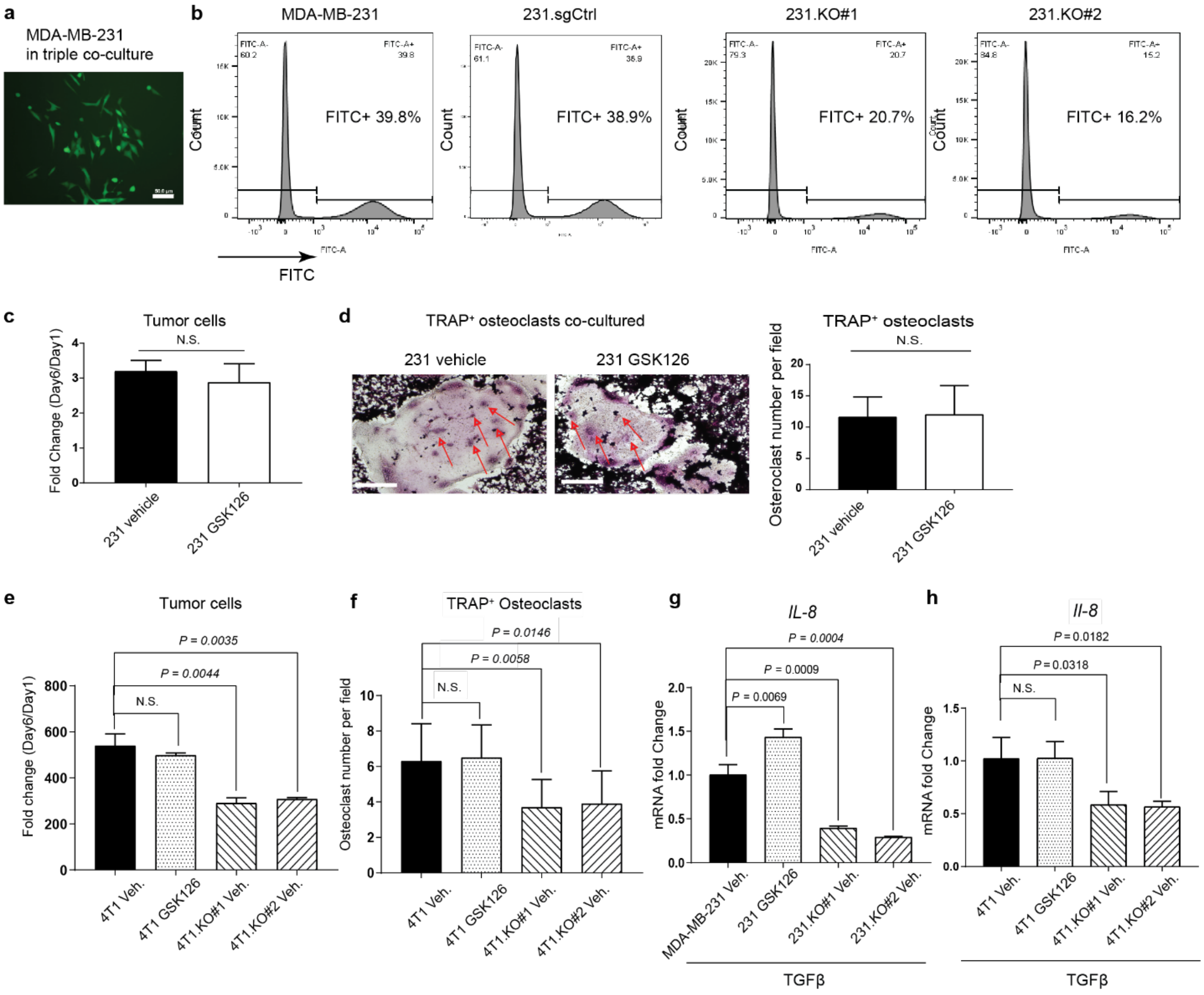


**Supplementary Fig. 2.** EZH2 enhances tumor cell proliferation and osteoclast maturation in the triple co-culture system. **a,** Fluorescence image of MDA-MB-231 cells with expression of GFP in triple co-culture. Scale bars, 50 μm. **b,** Flow cytometric analysis of $GFP^+$ breast cancer cells isolated from triple co-culture as measured in the fluorescein isothiocyanate (FITC) channel. **c,** Quantification of MDA-MB-231 cells treated with a vehicle or GSK126 (2 μM) after co-culture with osteoclasts and MC3T3 osteoblasts and treated with TGFβ (5 ng/mL) for 6 days. Data are presented as means ± S.D. (*t*-test). N.S., not significant. **d,** Representative staining images and quantification of mature $TRAP^+$ osteoclasts after co-culture with MC3T3 osteoblasts and MDA-

MB-231 cells treated with a vehicle or GSK126 (2 μM), and TGFβ (5 ng/mL) for 6 days. Scale bars, 200 μm. Data are presented as means ± S.D. (*t*-test). **e,** Quantification of 4T1 cells treated with vehicle, or with GSK126 (6 μM), 4T1.KO#1 cells, and 4T1.KO#2 cells treated with vehicle, after co-culture with osteoclasts and MC3T3 osteoblasts and treated with TGFβ (5 ng/mL) for 6 days. Data are presented as means ± S.D. (*t*-test). **f,** Quantification of mature $TRAP^+$ osteoclasts after co-culture with MC3T3 osteoblasts and the indicated cancer cells treated with TGFβ (5 ng/mL) for 6 days. Data are presented as means ± S.D. (*t*-test). **g,** qRT-PCR analysis of *IL-8* mRNA expression in the indicated cells treated with TGFβ (5 ng/mL, 2 hours). Data are presented as means ± S.D. (*t*-test). **h,** qRT-PCR analysis of *Il-8* mRNA expression in the indicated cells treated with TGFβ (5 ng/mL, 2 hours). Data are presented as means ± S.D. (*t*-test).

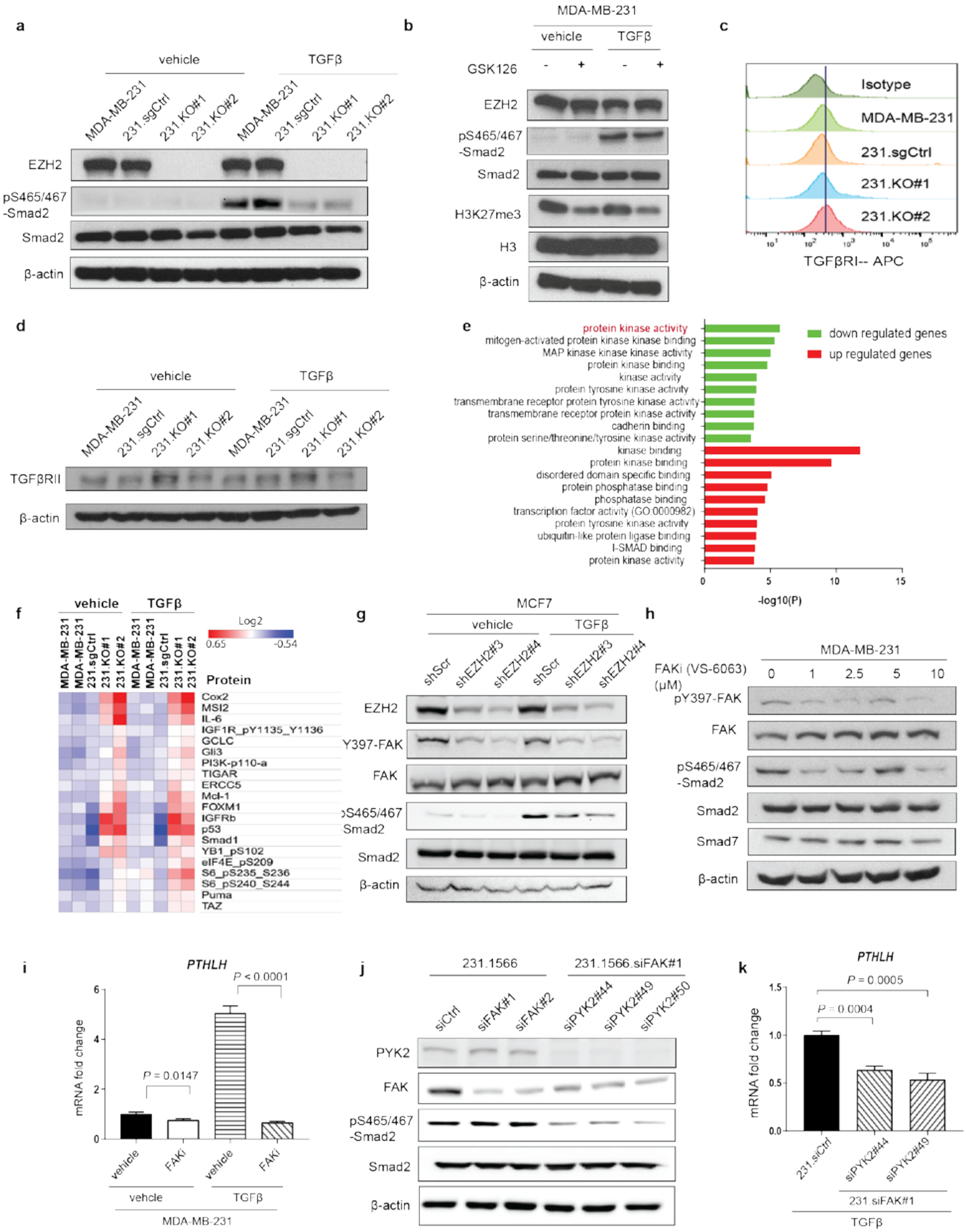


**Supplementary Fig. 3.** EZH2 regulates pY397-FAK level, which enhances pS465/467-Smad2

expression in response to TGFβ stimulation. **a,** Western blotting of the expression of the indicated proteins in MDA-MB-231, 231.sgCtrl, 231.KO#1, and 231.KO#2 cells treated with a vehicle or TGFβ (5 ng/mL) for 2 hours. **b,** Western blotting of the expression of the indicated proteins in MDA-MB-231 cells treated with a vehicle or GSK126 (2 μM, 24 hours) and then a vehicle or TGFβ (5 ng/mL, 2 hours). **c,** Results of flow cytometric analysis of TGFβRI in the indicated cancer cells as measured in the APC channel. **d,** Western blotting of the expression of TGFβRII and β-actin in the indicated cells treated with a vehicle or TGFβ (5 ng/mL, 2 hours). **e,** Gene Ontology (GO) molecular functional analysis of RPPA data showing the function of upregulated and downregulated proteins in 231.KO#1 and 231.KO#2 cells compared with MDA-MB-231 and 231.sgCtrl cells. **f,** Results of RPPA analysis of MDA-MB-231, 231.sgCtrl, 231.KO#1, and 231.KO#2 cells treated with a vehicle or TGFβ (5 ng/mL, 2 hours). The heat map shows the top upregulated proteins in 231.KO#1 and 231.KO#2 cells compared with MDA-MB-231 and 231.sgCtrl cells. **g,** Western blot of the expression of the indicated proteins in MCF7.shScr, MCF7.shEZH2#3, and MCF7.shEZH2#4 cells treated with a vehicle or TGFβ (5 ng/mL, 2 hours). **h,** Western blotting of the expression of the indicated proteins in MDA-MB-231 cells treated with the FAKi VS-6036 at different concentrations (0-10 μM) and with TGFβ (5 ng/mL, 2 hours). **i,** Results of qRT-PCR analysis of *PTHLH* mRNA expression in MDA-MB-231 cells treated with a vehicle or VS-6036 (10 μM, 24 hours) and then a vehicle or TGFβ (5 ng/mL, 2 hours). Data are presented as means ± S.D. (*t*-test). **j,** Western blotting of the expression of PYK2, FAK, pS465/467-Smad2, Smad2, and β-actin in the indicated 231-1566 cell sublines treated with TGFβ (5 ng/mL, 2 hours). **k,** Results of qRT-PCR analysis of *PTHLH* mRNA expression in 231.siCtrl, 231.siFAK#1 siPYK2#44, and 231.siFAK#1 siPYK2#49 cells treated with TGFβ (5 ng/mL, 2 hours). Data are presented as means ± S.D. (*t*-test).

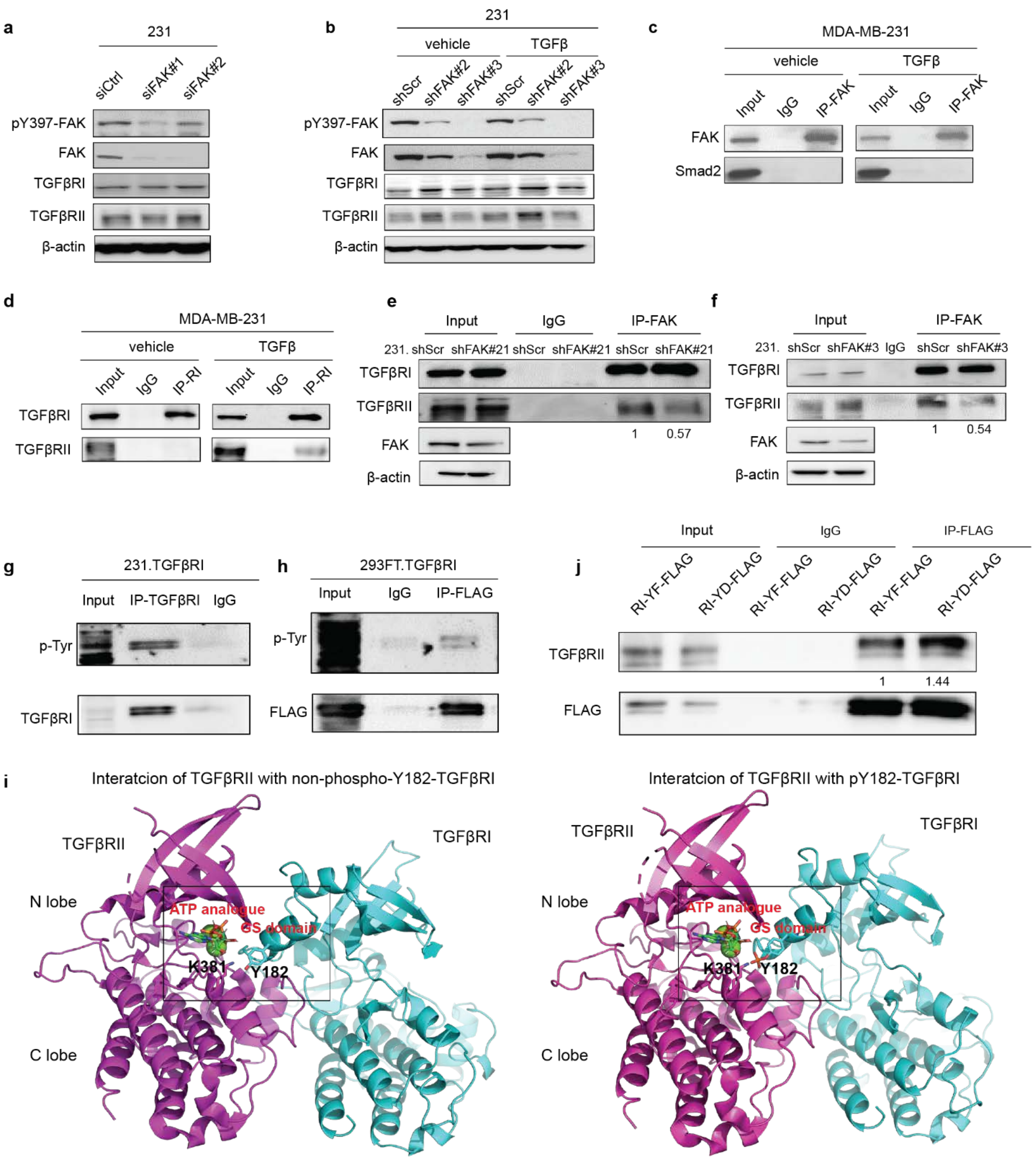


**Supplementary Fig. 4.** FAK inhibitor blocks TGFβRI binding to TGFβRII. **a,** Western blotting of the expression of the indicated proteins in 231.siCtrl, 231.siFAK#1, and 231.siFAK#2 cells. **b,** Western blotting of the expression of the indicated proteins in 231.shCtrl, 231.shFAK#2, and 231.siFAK#3 cells treated with a vehicle or TGFβ (5 ng/mL, 2 hours). **c,** IP of FAK from the lysis of MDA-MB-231 cells treated with a vehicle or TGFβ (5 ng/mL, 2 hours) followed by Western blotting for FAK and Smad2. **d,** IP of TGFβRI from the lysis of MDA-MB-231 cells treated with a vehicle or TGFβ (5 ng/mL, 2 hours) followed by Western blotting for TGFβRI and TGFβRII. **e,** IP of FAK from 231.shScr, and 231.shFAK#21 cell lysis followed by Western blotting for TGFβRI and TGFβRII. Western blotting detected FAK and β-actin in the inputs of 231.shScr and 231.shFAK#21 cells. **f,** IP of FAK from 231.shScr and 231.shFAK#3 cell lysis followed by Western blotting for TGFβRI and TGFβRII. Western blotting detected FAK and β-actin in the inputs of 231.shScr and 231.shFAK#3 cells. **g**, MDA-MB-231 cells were transfected with FLAG tagged TGFβRI plasmid (231.TGFβRI). IP of FLAG from the lysis of 231.TGFβRI cells using anti-FLAG antibody, followed by Western blotting for phospho-Tyrosine. **h,** HEK293FT cells were transfected with FLAG tagged TGFβRI plasmid (293FT.TGFβRI). IP of FLAG from the lysis of 293FT.TGFβRI cells using anti-FLAG antibody, followed by Western blotting for phospho-Tyrosine. **i,** Docked complex models showing the interaction between GS domain of TGFβRI (without/with phosphorylation of Y182) and catalytic site of TGFβRII. **j,** HEK293FT cells were transfected with TGFβRII and FLAG tagged TGFβRI-Y182F mutant (RI-YF-FLAG) or TGFβRI-Y182D mutant (RI-YD-FLAG). IP of FLAG from the lysis of cells using anti-FLAG antibody, followed by Western blotting for TGFβRII.

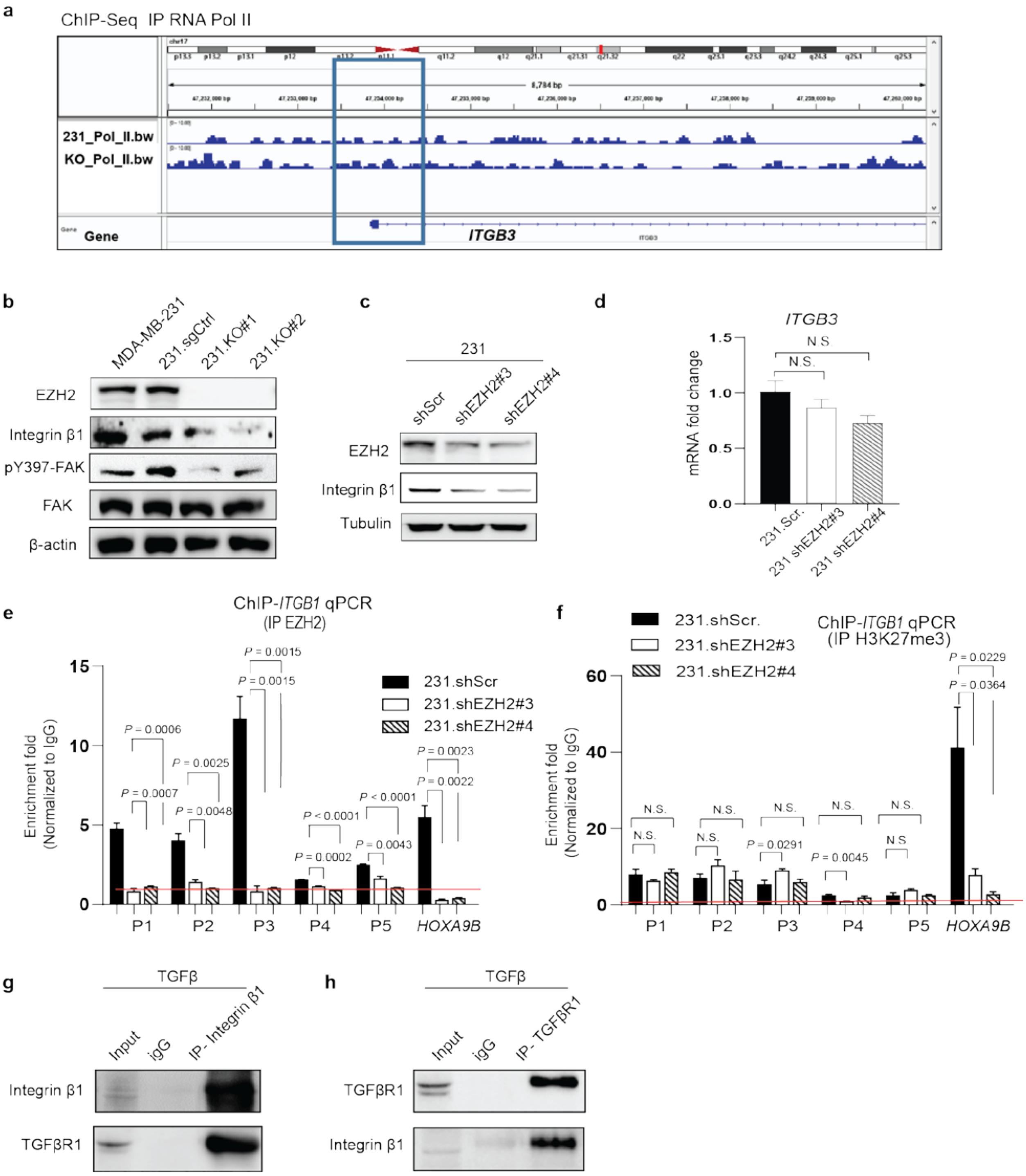


**Supplementary Fig. 5.** EZH2 increases *ITGB1* mRNA expression and binds at the promoter of *ITGB1* gene. **a,** Screenshot of the RNA Pol II ChIP sequencing (ChIP-seq) signal at the *ITGB3 promoter* locus in MDA-MB-231 (231_Pol _II.bw) and 231.KO#1 (KO_Pol_I.bw) cells. **b,** Western blotting of the expression of the indicated proteins in MDA-MB-231, 231.sgCtrl,

231.KO#1, and 231.KO#2 cells. **c,** Western blotting of the expression of the indicated proteins in 231.shSrc, 231.shEZH2#3, and 231.shEZH2#4 cells. **d,** Results of qRT-PCR analysis of *ITGB3* mRNA expression in the indicated cells. Data are presented as means ± S.E.M. (*t*-test). **e,** EZH2 was immunoprecipitated from 231.shSrc, 231.shEZH2#3, and 231.shEZH2#4 cells, and EZH2 binding to *ITGB1* was detected using qPCR with the indicated primers. *HOXA9B* was used as a positive control, and all fold-enrichment values were normalized according to IgG values. Data are presented as means ± S.E.M. (*t*-test). **f,** H3K27me3 was immunoprecipitated from 231.shSrc, 231.shEZH2#3, and 231.shEZH2#4 cells, and H3K27me3 binding to *ITGB1* or *HOXA9B* was detected using qPCR with the indicated primers. *HOXA9B* was used as a positive control, and all fold-enrichment values were normalized according to IgG values. Data are presented as means ± S.E.M. (*t*-test). **g,** Integrin β1 protein was immunoprecipitated from the lysis of MDA-MB-231 cell treated with a vehicle or TGFβ (5 ng/mL, 2 hours) by anti-integrin β1 antibodies, which was followed by Western blotting for TGFβRI. **h,** TGFβRI protein was immunoprecipitated from the lysis of MDA-MB-231 cell treated with a vehicle or TGFβ (5 ng/mL, 2 hours) by TGFβRI antibodies, which was followed by Western blotting for Integrin β1.

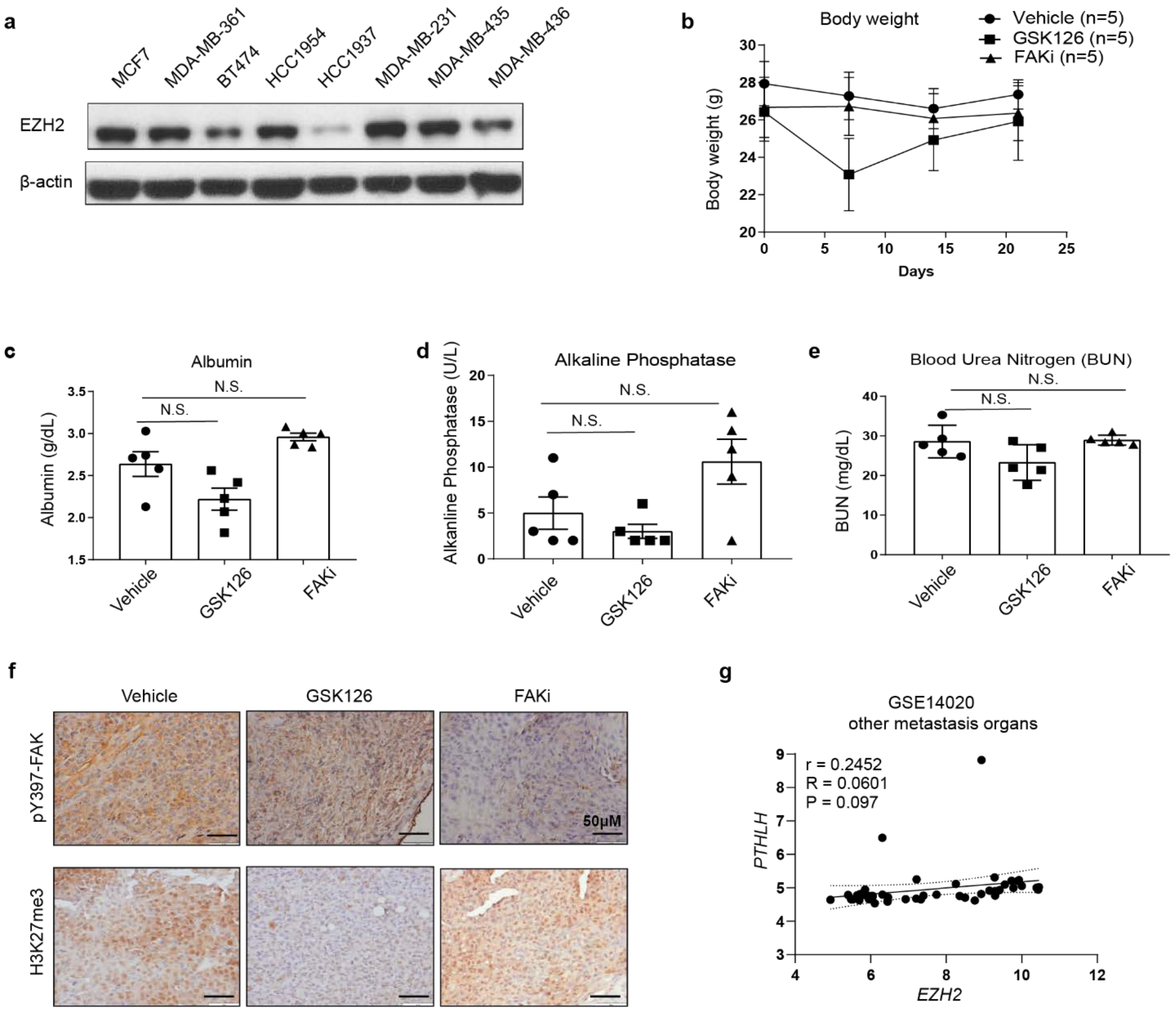


**Supplementary Fig. 6.** FAK inhibitor VS-6063 blocks breast cancer bone metastasis without side effect. **a,** Western blotting of the expression of EZH2 and β-actin in the indicated breast cancer cells. **b,** Body-weight curves for three subgroups of mice intratibially injected with MDA-MB-231 cells and given treatment with a vehicle, GSK126 (100 mg/kg/day, i.p. injection), or the FAKi VS-6063 (50 mg/kg, twice a day, oval gavage) beginning on day 18 after injection. Data are presented as means ± S.D. **c-e,** The albumin (**c**), alkaline phosphatase (**d**), and blood urea nitrogen (**e**) expression levels in the blood of the three subgroups of mice in **b**. Data are presented as means ± S.D. (*t*-test). N.S., not significant. **f,** Representative pictures of IHC staining of pY397-FAK and

H3K27me3 expression in the bone metastasis samples obtained from the three subgroups of mice in **b**. **g,** The Pearson *r* correlation for *EZH2* and *PTHLH* RNA mRNA expression in lung, liver, and brain metastases in breast cancer patients (GSE14020 data set).
